# Improving vocal communication with a ketogenic diet in a mouse model of autism

**DOI:** 10.1101/2023.10.05.561083

**Authors:** Dorit Möhrle, Kartikeya Murari, Jong M Rho, Ning Cheng

## Abstract

**Background:** Deficits in social communication and language development is a hallmark of autism spectrum disorder currently with no cure. Interventional studies using animal models have been very limited in demonstrating improved vocal communication. Autism has been proposed to involve metabolic dysregulation. Ketogenic diet (KD) is a metabolism-based therapy for medically intractable epilepsy, and its applications in other neurological conditions have been increasingly tested. However, how it would affect vocal communication has not been explored. The BTBR mouse strain is considered a model of idiopathic autism. They display robust deficits in vocalization during social interaction, and have metabolic changes implicated in autism.

**Methods:** We investigated the effects of KD on ultrasonic vocalizations (USVs) in juvenile and adult BTBR mice during male-female social encounters.

**Results:** After a brief treatment with KD, the amount, spectral bandwidth, and much of the temporal structure of USVs were robustly improved in both juvenile and adult BTBR mice. Composition of call categories and transitioning between individual call subtypes was more effectively improved in juvenile BTBR mice.

**Limitations:** Although sharing certain attributes, mouse vocalization is unlikely to model all aspects in the development and deficits of human language. KD is highly restrictive and can be difficult to administer, especially for many people with autism who have narrow food selections. Side effects and potential influence on development should also be considered. Future studies are required to tease apart the molecular mechanisms of KD’s effects on vocalization.

**Conclusions:** Together, our data provide further support to the hypothesis that metabolism-based dietary intervention could modify disease expression, including core symptoms, in autism.

## 1. Introduction

Autism spectrum disorder is a prevalent neurodevelopmental disorder, characterized by deficits in social function and language development, as well as repetitive and restrictive behaviours (DiCicco-Bloom et al., 2006; Geschwind 2009; Lai et al., 2014; Llaneza et al., 2010). Currently, there is no effective treatment for the core symptoms.

Language impairment is central to autism (Stefanatos and Baron 2011; Krishnan et al., 2016; Mayes et al., 2015; Rapin and Dunn 2003; Brignell et al., 2018). Currently, behavioral intervention is the principal treatment modality, but has yielded limited results (Spreckley and Boyd 2009; Paul 2008). It also requires extensive training and ongoing support from parents, teachers, and paraprofessionals. Studies focused on deficits in vocal communication have been conducted in animal models, including mice (Lahvis et al., 2011; Konopka and Roberts 2016; Fischer and Hammerschmidt 2011). Ultrasonic vocalizations (USV) in mice are remarkably complex, biologically meaningful signals (Lahvis et al., 2011; Arriaga and Jarvis 2013; Portfors and Perkel 2014), with relevant parameter including not only the amount of vocalizations, but also the acoustic features, structures of call organization, and sonographic categories (Takahashi et al., 2016; Castellucci et al., 2018; Panksepp et al., 2007; Wang et al., 2008; Holy and Guo 2005). Many mouse models of autism display impaired USV, including ones that model genetic alterations such as Shank1/2/3, neuroligin-3/4, Tsc2, Fmr1, 15q11-13, 16p11.2 (Jamain et al., 2008; Young et al., 2010; Ju et al., 2014; Wöhr 2014; Kogan et al., 2015; Yang et al., 2015; Brunner et al., 2015; Stoppel et al., 2018; Rotschafer et al., 2012), as well as ones mimicking environmental risks such as maternal immune activation and exposure to valproic acid (Malkova et al., 2012; Cezar et al., 2018). In addition, autism risk genes involved in human speech and language development such as *Foxp2*, *Foxp1* and *Cntnap2* also play a role in mouse vocalization, supported by findings in respective transgenic mouse models that display altered USV behavior (Peñagarikano et al., 2011; Brunner et al., 2015; Castellucci et al., 2016; Chabout et al., 2016; Schaafsma et al., 2017; Usui et al., 2017). However, unlike other core autism-like traits such as social approach and repetitive behavior that can be improved by diverse pharmacological treatments in a multitude of animal models (Kazdoba et al., 2016), very limited reports of improving vocalization exist (Rotschafer et al., 2012; Cezar et al., 2018). In these studies, vocalization amount, but not other acoustic features, was altered in the animal models and rescued by pharmacological treatments.

Autism has been proposed a disorder involving metabolic dysregulation (Rossignol and Frye 2012; Cheng et al., 2017; Napoli et al., 2014; Mayengbam et al., 2021), and a metabolic therapy in use for nearly a century is the ketogenic diet (KD). Characterized by a high-fat and low-carbohydrate content, KD is a proven therapy for treating medically intractable epilepsy (Neal et al., 2008). Recently, it has been tested in a wider variety of neurological diseases including Alzheimer’s disease, Parkinson’s disease, amyotrophic lateral sclerosis, sleep disorders, multiple sclerosis, brain trauma, stroke, pain, Huntington’s disease and brain cancer (Cheng et al., 2017; Stafstrom and Rho 2012; Boison 2016; Gano et al., 2014; Lutas and Yellen 2013; Gasior et al., 2006; Yudkoff et al., 2007). In addition, several studies (although limited in scope) have shown promising results in some patients with autism (Evangeliou et al., 2003; Herbert and Buckley 2013; Spilioti et al., 2013; Lee et al., 2018). In addition, KD reduces deficits in social interaction and repetitive behaviors, as well as comorbid conditions such as seizures, in several rodent models of autism (Ruskin et al., 2017a; Ruskin et al., 2017b; Ruskin et al., 2013; Nylen et al., 2008; Ahn et al., 2014; Westmark et al., 2020). However, its effect on vocalization has not been explored.

The BTBR T+tf/J (BTBR) inbred mouse strain has been well characterized with various tests for social interaction and communication including USV (Scattoni et al., 2008; Scattoni et al., 2011), as well as for repetitive behaviours, traits that model the defining symptoms of autism (for review see e.g.Möhrle et al., 2020). In general, BTBR mice display robust phenotype in these assays (Meyza and Blanchard 2017; Meyza et al., 2013). In addition, a relevant anatomical feature of BTBR mice is agenesis of the corpus callosum (Meyza and Blanchard 2017), a structure in which alterations have been consistently found in human studies (Frazier and Hardan 2009; Wolff et al., 2015; Paul et al., 2014). Thus, BTBR mouse has been considered a valuable model of idiopathic autism and used to test many potential therapies, which provided the scientific rationale for a multitude of clinical trials (Kazdoba et al., 2016). Relevant to this study, previous reports have pointed to links between metabolism and behavior in BTBR mice through neuroinflammation, neurogenesis, transcriptome/proteome and microbiota in manners that reflect autism in humans (Flowers et al., 2007; Stern 2011; Currais et al., 2016; Golubeva et al., 2017; Cheng et al., 2017; Rivell and Mattson 2019; Daimon et al., 2015).

Here, to test how KD affect vocalization, we fed juvenile and adult BTBR mice either standard diet (SD) or KD, and quantified their USV during male-female interaction. We also investigated C57BL/6J (B6) mice, a strain with normal levels of social behaviors (Meyza and Blanchard 2017), including vocalization (Scattoni et al., 2008; Scattoni et al., 2011; Wöhr et al., 2011; Scattoni et al., 2013; Schwartzer et al., 2013; Wöhr 2015; Kim et al., 2016). Our results demonstrated that after a brief treatment with KD, the amount, spectral bandwidth, and many aspects of the temporal profile of USVs in BTBR mice were robustly improved in both juvenile and adult stage. KD treatment was more effective to improve the composition of call type categories in juvenile than adult BTBR mice. Finally, KD normalized transitions between call types in juvenile, but not adult, BTBR mice. Fewer changes through KD diet occurred in B6 mice.

## 2. Materials and Methods

### 2.1. Mice

Breeder B6 and BTBR animals were obtained from the Jackson Laboratory (ME) and the lines were maintained at the mouse facility of the Cumming School of Medicine, University of Calgary. Mice were group-housed with up to five mice per cage in a humidity-and temperature-controlled room with a 12-h light/dark cycle and were fed *ad libitum*. Both juvenile and adult B6 and BTBR male animals were fed either standard mouse chow (standard diet, SD) or KD. Sexually mature female B6 and BTBR mice 8-10 weeks of age were used in USV tests. All procedures in this study were performed in accordance with the recommendations in the Canadian Council for Animal Care. The protocol of this study was approved by the Health Sciences Animal Care Committee of the University of Calgary.

### 2.2. Administration of KD

Male juvenile or adult B6 and BTBR mice were randomly assigned to two groups: 1) SD (5062, Lab Supply, TX); 2) KD (weight ratio of fat to carbohydrate plus protein is approximately 6.3:1, F3666, Bio-serv, NJ). Percentage calories provided by each nutrient category are: for SD, 23.2% form protein, 21.6% from fat, and 55.2% from carbohydrates; for KD, 4.7% form protein, 93.4% from fat, and from 1.8% carbohydrates. Detailed nutritional information of the diets is included in Supplementary Material (Supplementary Figure 1, 2). All groups were fed *ad libitum*. For juvenile mice, diet treatment began around postnatal day 30, and lasted 11 days until animals were sacrificed. For adult mice, dietary treatment began around 8 weeks of age and also lasted 11 days. Together, there were four experimental groups tested at each age: B6_SD, B6_KD, BTBR_SD, and BTBR_KD.

### 2.2. USV recording apparatus and procedure

SD-or KD-fed mice were tested in parallel to minimize variation in experimental conditions. Male subject mice were housed individually in clean standard cages for 3-4 hours before testing. Most bedding was removed from the cage to reduce background noise. Tests were carried out between 6 to 8 pm (Scattoni et al., 2011). USVs were recorded with a CM16/CMPA condenser microphone positioned about 20 centimetres above the cage and connected to a UltraSoundGate 116H digitizer (Avisoft Bioacoustics, Berlin, Germany). Recordings were monitored with a computer running Avisoft RECORDER USGH software. A mature (8-10 week old) female mouse of the same strain was introduced into the home cage of the male subject for 5 min. The male subjects exhibited typical chasing and sniffing behaviors. Mounting did not occur routinely, and when it did, it usually happened toward the end of the 5 min testing period. USVs between 0-200 kHz were recorded during this dyadic male-female interaction over a 5 min period (Scattoni et al., 2011). Before the test, neither male nor female mice had been exposed to the opposite sex after weaning. Live female mice instead of female urine or anesthetized females were used to model natural dyadic social interactions. Previous studies have found that in this situation, the majority (∼80%) of USVs emitted is from the male mice (Warburton et al., 1989; Neunuebel et al., 2015), and males start to emit USVs around P19 (Warburton et al., 1989).

### 2.4. USV analysis

Audio files were analyzed offline using the VocalMat software suite (Fonseca et al., 2021) for MATLAB (version R2022a, The MathWorks, Inc., Natick, Massachusetts) to detect and classify USVs into one of 11 call types (step up, step down, chevron, reverse [rev] chevron, down frequency modulated, up fm, complex, flat, short, two steps, multi steps). Call type distribution as well as simple categorical and spectral USV features including total number of calls, duration of calls, mean frequency and bandwidth were analyzed from VocalMat call logs (exported to Excel 2016, Microsoft, Redmond, Washington) similar to Möhrle et al., (2023) using custom-written MATLAB scripts. As previously reported (Takahashi et al., 2016; Castellucci et al., 2018; Panksepp et al., 2007; Wang et al., 2008; Holy and Guo 2005), USVs were not evenly produced in time; instead, they were generated in clusters punctuated by periods of silence. This temporal organization of USV density was analyzed modified from Möhrle et al., (2023). In brief, two distinct inter-call intervals (ICIs) were defined based on previous findings (Hage et al., 2013; Chen et al., 2021; Castellucci et al., 2018) and characteristics of temporal USV density extracted from calls emitted in sequences (separated by ICIs ≥100 ms) and bouts (ICIs ≥2000 ms). For call type syntax analysis, the VocalMat Excel logs were reorganized and transition probabilities between call types within call bouts calculated and displayed in syntax flow paths using DeepSqueak (version 2.6.2, Coffey et al., 2019) for MATLAB. As for temporal organization analysis, an inter-bout interval of 2 s was chosen (maximum bout separation 2s, exclude classes with frequency below 0.01).

### 2.5. Statistical analysis

Statistical tests were performed in GraphPad Prism 9.5.1 (GraphPad Software, San Diego, California) and RStudio 2022.2.0.0 (PBC, Boston, Massachusetts), and Figures were generated in GraphPad Prism and in DeepSqueak for MATLAB (Coffey et al., 2019). Data following normal distribution (D’Agostino & Pearson test) were analyzed using two-way ANOVA and *post hoc* multiple comparison tests with correction for type 1 error after Tukey’s method. To accommodate for non-normal distribution or data normalization we used ARTool (Aligned Rank Transform, ART) to align-and-rank data for nonparametric two-way or repeated measures ANOVA for main effects and interactions (Wobbrock et al., 2011), and ART-C for *post hoc* pairwise comparisons (contrast tests, Elkin et al., 2021). All statistical analyses are presented in the Figure legends or respective Tables. Statistical significance level was *α* = 0.05, and resulting *p* values are reported in the legends and Tables using: \**p* < 0.05; \*\**p* < 0.01; *p*\*\*\* < 0.001. Unless stated otherwise, normally distributed data are presented as group mean, standard deviation (StdDev) and individual data points (animals); non-normally distributed data as group median, interquartile range (IQR) and individual data points (animals). Syntax flow paths are descriptive.

## 3. Results

### 3.1. KD reversed deficits in the amount and bandwidth of USV in both juvenile and adult BTBR mice

In a previous study, we explored structural, metabolic, and functional consequences of a brief treatment of a KD (weight ratio of fat to carbohydrate plus protein is approximately 6.3:1) in naive juvenile mice of different inbred strains, including the BTBR mice (Mayengbam et al., 2021). Here, to determine the effect of KD on vocalization, we recorded USVs in both B6 and BTBR mice during male-female interaction (Scattoni et al., 2011) after mice were fed SD or KD. Consistent with previous findings (Scattoni et al., 2011), both juvenile and adult BTBR mice emitted significantly less USVs than B6 mice, with total calls being only ∼40% of those from B6 (Fig. 1A, B, Table 1). In addition, the mean frequency and bandwidth was lower in both juvenile and adult BTBR mice (Fig. 1E-H, Table 1), whereas the duration of individual calls did not differ between strains (Fig. 1C, D, Table 1). A brief treatment with KD for 11 days robustly increased the call amount in both juveniles and adults independent of strain (Fig 1A, B, Table 1). In addition, in juveniles, KD increased the bandwidth of calls from BTBR mice to a similar level as B6 mice (Fig. 1G, Table 1, 2), whereas in adults both KD-fed BTBR and B6 mice emitted calls with a higher bandwidth (Fig. 1H, Table 1). In contrast, KD did not rectify the lower mean frequency in BTBR mice from either age group (Fig. 1E, F, Table 1, 2), and left the duration of individual calls largely unchanged (Fig. 1C, D, Table 1, 2).

**Fig. 1.**
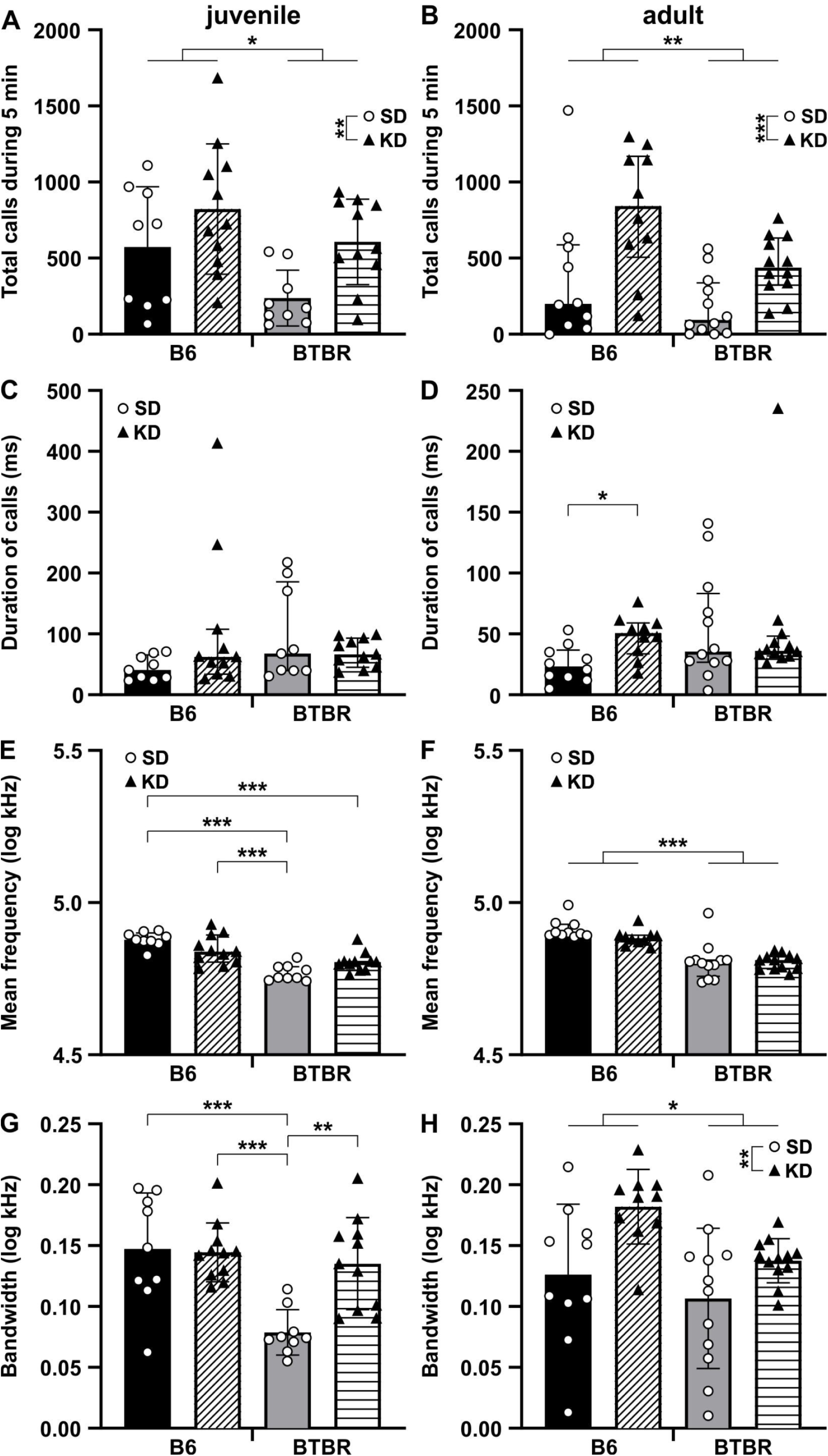
KD robustly reversed the deficits in the amount and spectral bandwidth features of USVs in juvenile and adult BTBR mice. **(A-H)** Quantification of total number of calls in 5-minute recordings, as well as duration, mean frequency and bandwidth of calls in juvenile **(A, C, E, G)** and adult **(B, D, F, H)** BTBR or B6 mice fed with either SD (circles) or KD (triangles). (**A, B**) The total number of calls was lower in BTBR mice. KD diet increased the number of calls independent of strain. **(C, D)** Average duration of individual calls was similar between strains. KD led to longer call duration only in (**D**) adult B6_KD. Note that the Y axes are different between juveniles and adults. (**E**) The mean frequency of calls was lower in juvenile BTBR_SD and BTBR_KD mice compared with B6_SD. (**F**) KD had no effect on the lower mean frequency in adult BTBR mice. (**G**) Call bandwidth was significantly lower in BTBR_SD compared with B6_SD mice. In BTBR_KD mice, the call bandwidth was higher than in BTBR_SD and similar to B6_SD. (**H**) Adult BTBR mice showed lower call bandwidth than B6 mice. KD increased the bandwidth independent of strain. Juveniles, B6_SD (*n*=9), B6_KD (*n*=11), BTBR_SD (*n*=9), and BTBR_KD (*n*=11); adults, B6_SD (*n*=10), B6_KD (*n*=10), BTBR_SD (*n*=12), and BTBR_KD (*n*=12). Data expressed as **(A, G, H)** mean (bars) ± StdDev (error bars) and individual animals (symbols) or **(B-F)** median (bars) ± IQR (error bars) and individual animals (symbols). *p* values, *** *p* < 0.0001, ** *p* < 0.01 and * *p* < 0.05.

**Table 1.**
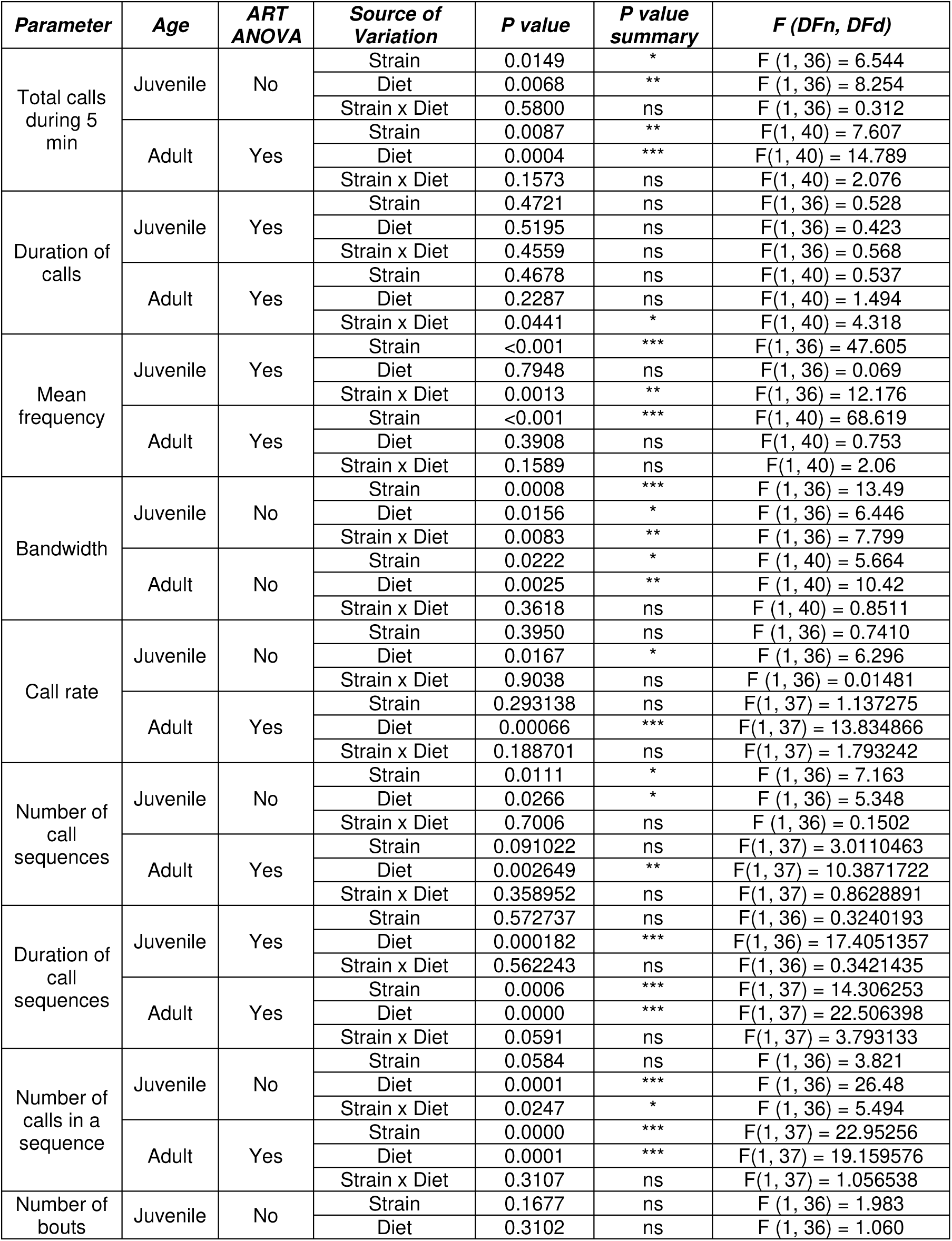
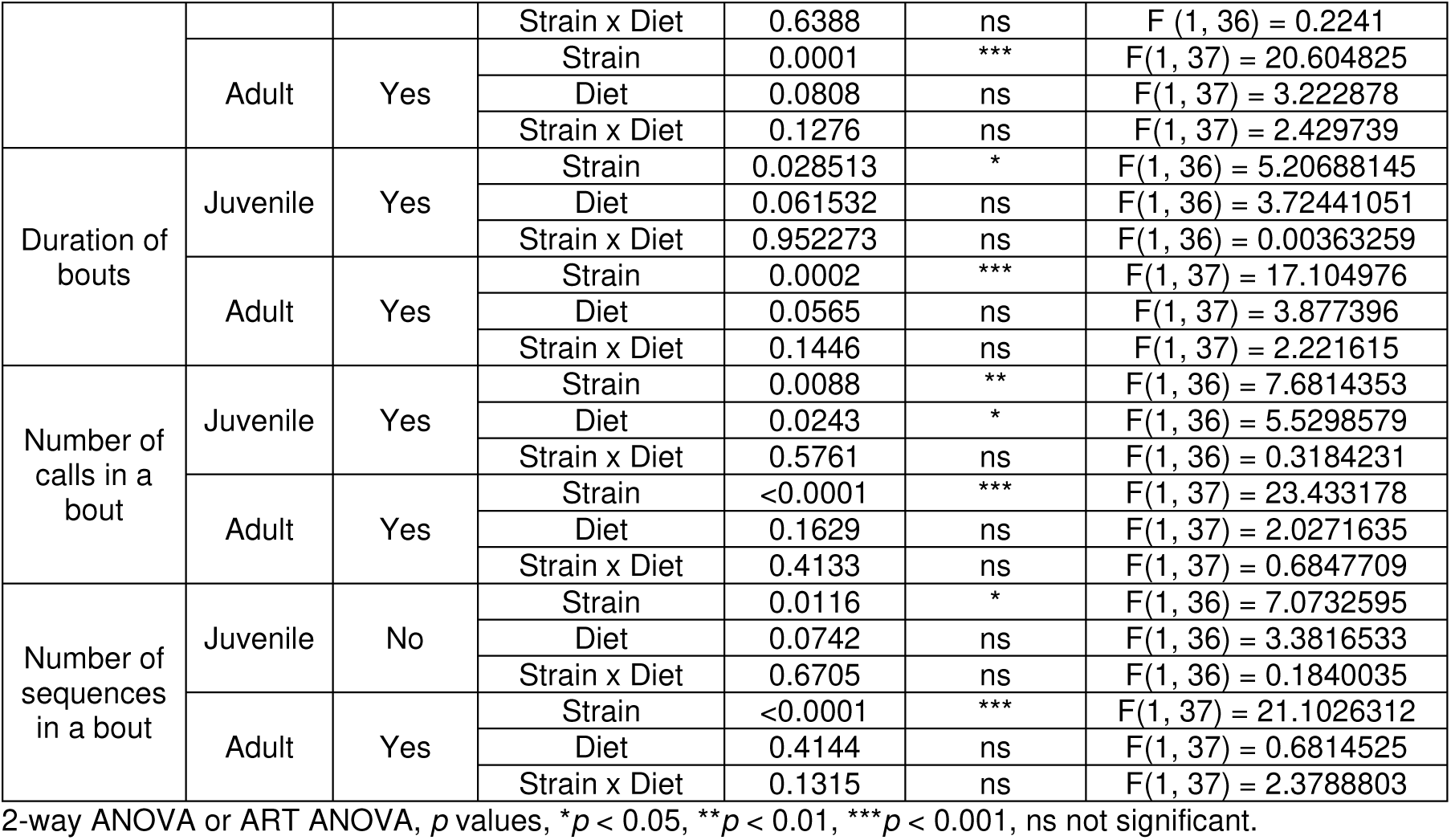
Statistical comparisons of USV acoustic and temporal features of calls from mice of the following four groups: juveniles, B6_SD (n=9), B6_KD (n=11), BTBR_SD (n=9), and BTBR_KD (n=11); adults, B6_SD (n=10), B6_KD (n=10), BTBR_SD (n=12), and BTBR_KD (n=12).

**Table 2.** *Post hoc* pairwise comparisons for Strain x Diet interactions of USV acoustic and temporal features of calls from mice of the following four groups: juveniles, B6_SD (n=9), B6_KD (n=11), BTBR_SD (n=9), and BTBR_KD (n=11); adults, B6_SD (n=10), B6_KD (n=10), BTBR_SD (n=12), and BTBR_KD (n=12).

| Parameter | Age | ART ANOVA | Contrast | Estimate | SE | DF | t | Adjusted P Value | P value summary |
| --- | --- | --- | --- | --- | --- | --- | --- | --- | --- |
| Duration of calls | Adult | Yes | BTBR,KD - BTBR,SD | 0.4 | 4.896 | 40 | 0.085 | 0.9997 | ns |
|  |  |  | BTBR,KD - B6,KD | -3.5 | 5.135 | 40 | -0.681 | 0.9035 | ns |
|  |  |  | BTBR,KD - B6,SD | 11.8 | 5.135 | 40 | 2.298 | 0.1156 | ns |
|  |  |  | BTBR,SD - B6,KD | -3.9 | 5.135 | 40 | -0.763 | 0.8706 | ns |
|  |  |  | BTBR,SD - B6,SD | 11.4 | 5.135 | 40 | 2.217 | 0.1361 | ns |
|  |  |  | B6,KD - B6,SD | 15.3 | 5.364 | 40 | 2.853 | 0.0331 | * |
| Mean frequency | Juvenile | Yes | BTBR,KD - BTBR,SD | 8.8 | 3.502 | 36 | 2.504 | 0.0763 | ns |
|  |  |  | BTBR,KD - B6,KD | -8.5 | 3.322 | 36 | -2.572 | 0.0657 | ns |
|  |  |  | BTBR,KD - B6,SD | -15.9 | 3.502 | 36 | -4.540 | 0.0003 | *** |
|  |  |  | BTBR,SD - B6,KD | -17.3 | 3.502 | 36 | -4.944 | 0.0001 | *** |
|  |  |  | BTBR,SD - B6,SD | -24.7 | 3.673 | 36 | -6.716 | <0.001 | *** |
|  |  |  | B6,KD - B6,SD | -7.4 | 3.502 | 36 | -2.100 | 0.1725 | ns |
| Bandwidth | Juvenile | No | BTBR,KD - BTBR,SD | 0.06 | 0.015 | 36 | 5.332 | 0.0031 | ** |
|  |  |  | BTBR,KD - B6,KD | -0.009 | 0.014 | 36 | 0.928 | 0.9128 | ns |
|  |  |  | BTBR,KD - B6,SD | -0.01 | 0.015 | 36 | 1.134 | 0.8532 | ns |
|  |  |  | BTBR,SD - B6,KD | 0.066 | 0.015 | 36 | 6.212 | 0.0005 | *** |
|  |  |  | BTBR,SD - B6,SD | -0.07 | 0.016 | 36 | 6.165 | 0.0006 | *** |
|  |  |  | B6,KD - B6,SD | -0.003 | 0.015 | 36 | 0.254 | 0.9979 | ns |
| Number of calls in a sequence | Juvenile | No | BTBR,KD - BTBR,SD | 2.0 | 0.374 | 36 | 7.49 | <0.0001 | *** |
|  |  |  | BTBR,KD - B6,KD | 0.1 | 0.355 | 36 | 0.410 | 0.9914 | ns |
|  |  |  | BTBR,KD - B6,SD | 0.8 | 0.374 | 36 | 3.191 | 0.1276 | ns |
|  |  |  | BTBR,SD - B6,KD | 1.9 | 0.374 | 36 | 7.101 | <0.0001 | *** |
|  |  |  | BTBR,SD - B6,SD | -1.1 | 0.392 | 36 | 4.099 | 0.0309 | * |
|  |  |  | B6,KD - B6,SD | 0.7 | 0.374 | 36 | 2.802 | 0.2139 | ns |

Tukey's or ART-C Tukey's multiple comparisons test, *p* values, \**p* < 0.05, \*\**p* < 0.01, \*\*\**p* < 0.001, ns not significant.

In summary, KD diet in both juvenile and adult BTBR mice resulted in more USV emission with increased call bandwidth, bringing these acoustic features closer to those of B6 mice, whereas the mean frequency was still lower.

### 3.2. KD normalized aspects of USV temporal structure quantified from call sequences

We further analyzed whether KD modulated the temporal USV structure, which is also a key element in vocal communication. As previously reported (Takahashi et al., 2016; Castellucci et al., 2018; Panksepp et al., 2007; Holy and Guo 2005; Wang et al., 2008), USVs were generated in clusters separated by periods of silence, instead of evenly distributed in time. To this end, we defined two distinct inter-call intervals (ICIs) thresholds: ICIs ≥100 ms separating sequences and ICIs ≥2000 ms separating bouts of calls (Fig. 2A). Our analysis of ICI length revealed a prominent peak at ∼60 to 80 ms for all groups (Fig. 2B), which was roughly consistent with the previously described ICI distribution (Hage et al., 2013). Thus, we chose 100 ms as the ICI threshold for separating call sequences. Additionally, 2-second pauses have been found as a natural threshold to define the end of a USV bout (Chen et al., 2021; Castellucci et al., 2018). Based on these ICI thresholds, we calculated features of call sequences and call bouts.

**Fig. 2.**
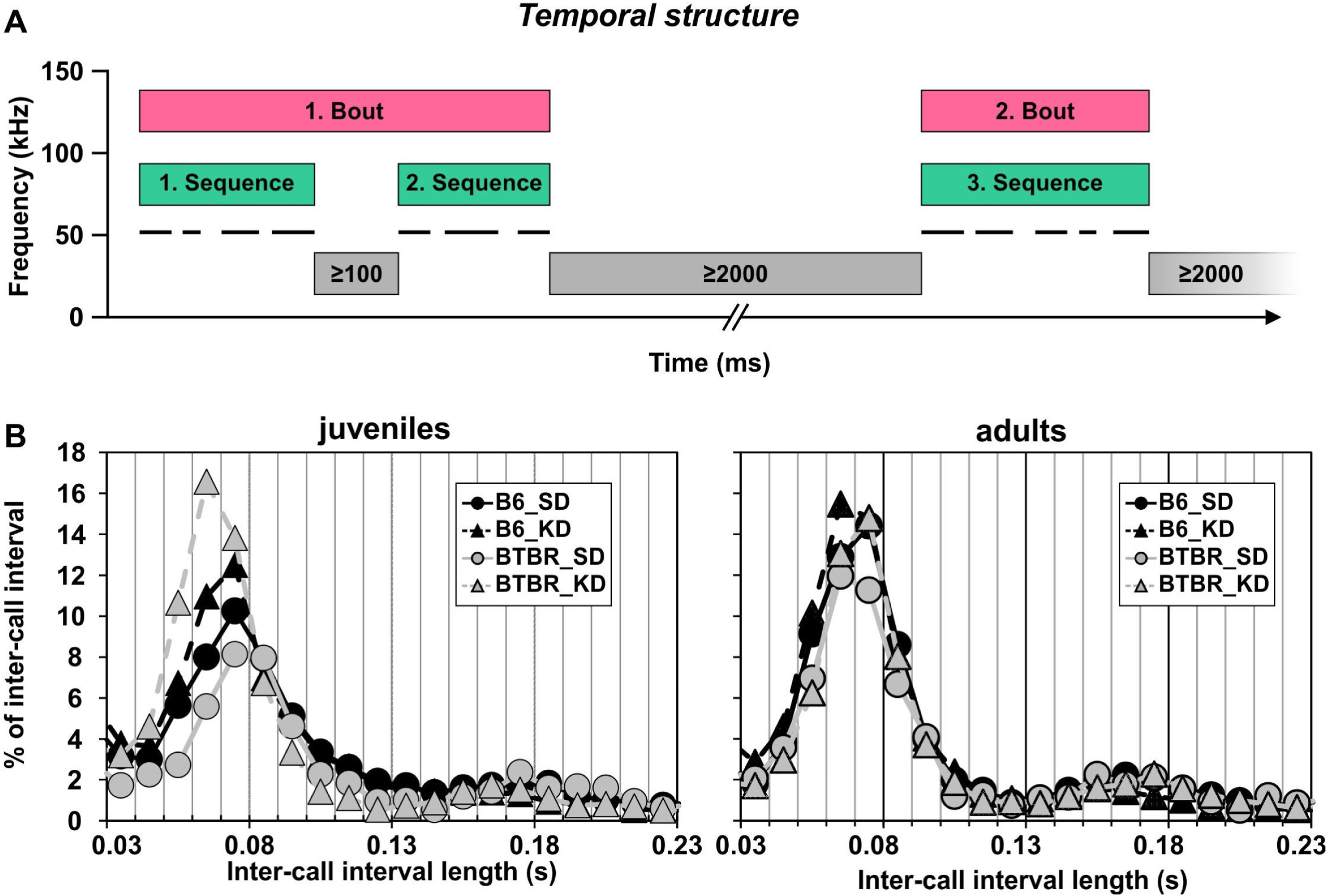
Temporal USV structure. **(A)** Schematic spectrogram of 50 kHz USVs (black horizontal lines). Two distinct inter-call interval (ICI) categories were defined as: ICIs ≥100 ms separating sequences of USVs, and ICIs ≥2000 ms separating bouts of USVs (adapted from Möhrle et al., 2023 © 2023 The Authors. Genes, Brain and Behavior published by International Behavioural and Neural Genetics Society and John Wiley & Sons Ltd.). (B) Histogram of ICI frequency distribution in juvenile (left) and adult (right) B6 and BTBR mice fed with either SD or KD diet. For all groups, there was a peak of ICIs falling into a bin around 60 to 70 and 70 to 80 ms. Juveniles, B6_SD (n=9), B6_KD (n=11), BTBR_SD (n=9), and BTBR_KD (n=11); adults, B6_SD (n=10), B6_KD (n=10), BTBR_SD (n=12), and BTBR_KD (n=12). Data expressed as relative frequency of numbers of ICIs (percentages) that lie within a bin. Bin width 0.01 s, bin centers 0.005 to 0.995s.

In juveniles, the total number of call sequences was reduced in BTBR mice independent of diet (Fig. 3A, Table 1), and the number of calls in a sequence was lower in SD-fed BTBR mice than in B6 mice (Fig. 3C, Table 1, 2). In contrast, the average duration of call sequences and the call rate within sequences (*i.e.,* number of calls per second) was similar between the two strains (Fig. 3E, G, Table 1). KD diet significantly increased the total number and duration of call sequences; and decreased the call rate within sequences irrespective of strain (Fig. 3A, E, G, Table 1). The total number of calls within a sequence was increased in BTBR_KD mice compared with BTBR_SD mice, and was similar to B6 mice of either diet (Fig. 3C, Table 1, 2).

**Fig. 3.**
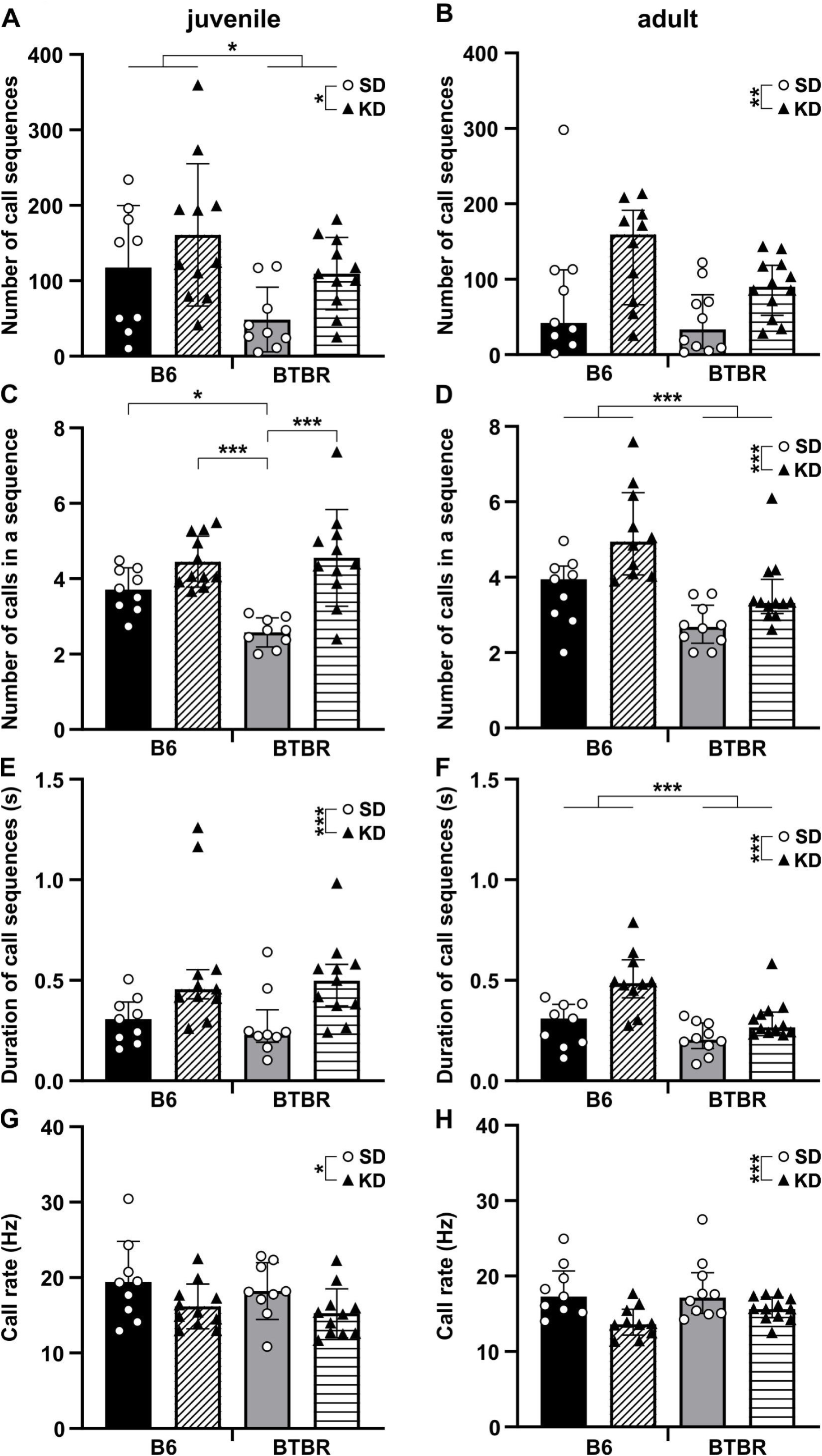
KD robustly reversed in deficits call sequence feature as measure of USV temporal structure in juvenile and adult BTBR mice. **(A-H)** Quantification of total number of call sequences in 5-minute recordings, as well as number of calls within a sequence, duration of call sequences and call rate within sequences in juvenile **(A, C, E, G)** and adult **(B, D, F, H)** BTBR or B6 mice fed with either SD (circles) or KD (triangles). (A) Total number of call sequences was lower in juvenile BTBR compared with B6 mice and increased through KD independent of strain. **(B)** In adults, the total number of call sequences was similar between strains and increased through KD in both B6 and BTBR mice. **(C)** Number of calls within a sequence was lower in BTBR_SD than in B6_SD and B6_KD mice. In BTBR_KD, the number of calls within a sequence was higher than in BTBR_SD and similar to either B6 group. **(D)** Number of calls in a sequence was decreased in BTBR mice and increased through KD in both B6 and BTBR mice. **(E)** Call sequences had similar duration in BTBR and B6 mice, and were longer after KD administration in both strains. **(F)** In adults, BTBR mice displayed shorter call sequences. KD prolonged the duration in both B6 and BTBR mice. **(G, H)** The call rate (calls per second within a sequence) was similar between strains and decreased through KD in **(G)** juvenile and **(H)** adult BTBR and B6 mice. Juveniles, B6_SD (*n*=9), B6_KD (*n*=11), BTBR_SD (*n*=9), and BTBR_KD (*n*=11); adults, B6_SD (*n*=10), B6_KD (*n*=10), BTBR_SD (*n*=12), and BTBR_KD (*n*=12). Data expressed as (C, G) mean (bars) ± StdDev (error bars) and individual animals (symbols) or (A, B, D-F, H) median (bars) ± IQR (error bars) and individual animals (symbols). *p* values, *** *p* < 0.0001, ** *p* < 0.01 and * *p* < 0.05.

In summary, juvenile BTBR mice emitted USVs organized in fewer numbers of call sequences, which in turn contained fewer calls. The KD diet normalized these aspects of temporal USV structure, but also induced a slower call emission and prolonged duration of sequences in both BTBR and B6 mice.

The USV temporal profile in adults varied slightly from that in juveniles. In adults, BTBR mice also displayed a lower number of calls within a sequence (Fig. 3B, Table 1), whereas the total number of call sequences was similar to B6 mice (Fig. 3D, Table 1). Additionally, the duration of call sequences was shorter in adult BTBR than B6 mice (Fig. 3F, Table 1). Similar to juveniles, the call rate was not different in adult BTBR compared with B6 mice (Fig. 3H, Table 1). The KD diet increased both the number of calls within a sequence and the duration of call sequences in BTBR and B6 mice (Fig. 3D, F, Table 1). Additionally, the total number of call sequences was higher (Fig. 3B, Table 1) and the call rate within a sequence lower (Fig. 3H, Table 1) in KD-fed than SD-fed BTBR and B6 mice.

In summary, adult BTBR mice emitted USVs in shorter call sequences with fewer calls in each sequence on average. The KD diet rectified both these parameters, but also led to a slower call emission organized in a greater total number of call sequences in both strains.

In a similar fashion, we quantified parameters of call bouts, *i.e.*, call successions with ICIs shorter than 2000 ms. Interestingly, the duration of bouts (Fig. 4A, B, Table 1), and the total number of calls (Fig. 4C, D, Table 1) or sequences within a bout (Fig. 4E, F, Table 1) were consistently lower in both juvenile and adult BTBR than B6 mice. However, KD only increased the number of calls within a bout in juvenile, but not adult, BTBR and B6 mice. The other aforementioned bout parameters remained unchanged through KD in either age group. The total number of bouts was similar in juvenile BTBR and B6 mice, and similar with SD and KD diet (Fig. 4G, Table 1). In contrast, adult BTBR mice showed a higher number of bouts than B6 mice which was not rectified through KD (Fig. 4H, Table 1).

**Fig. 4.**
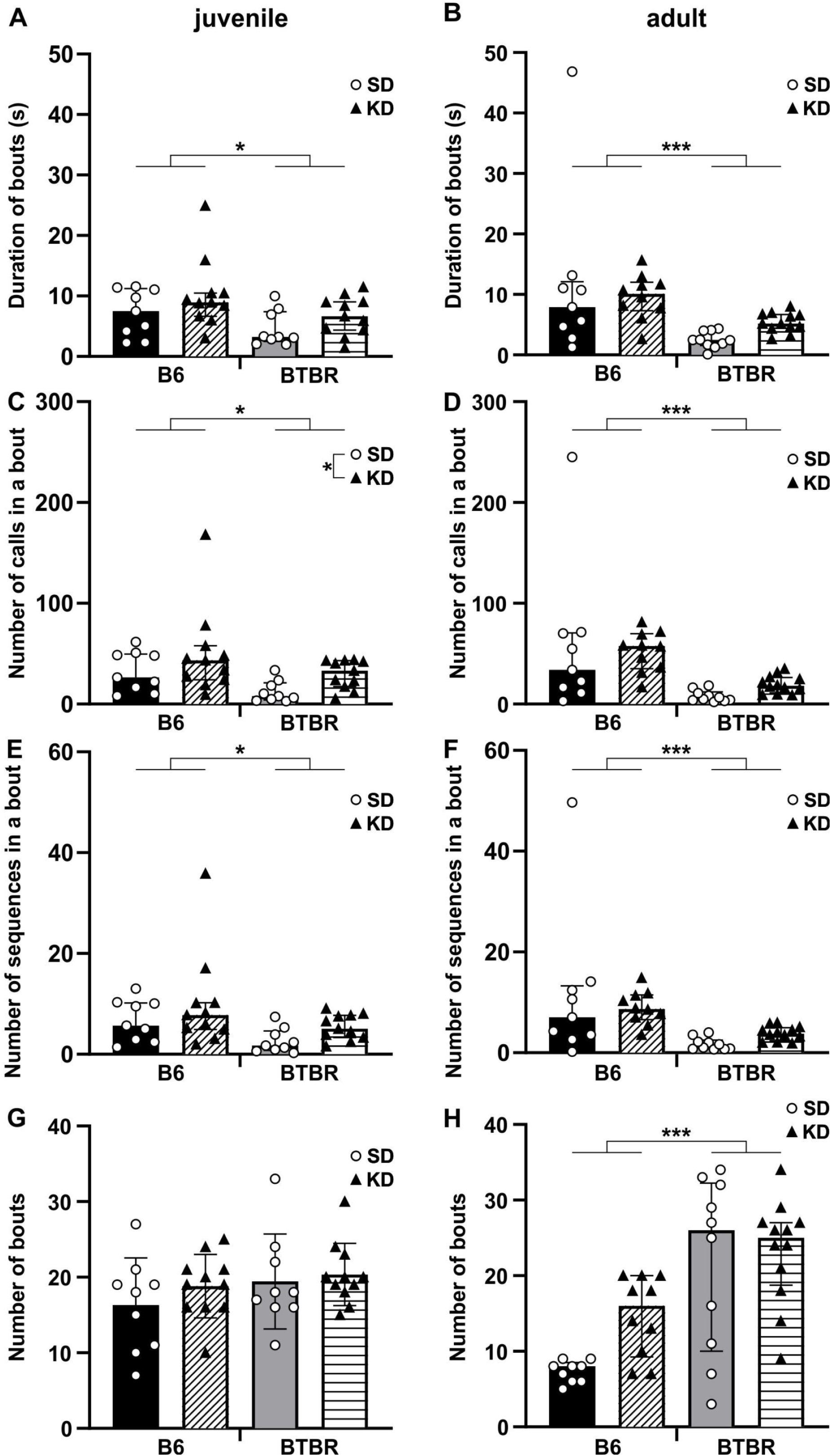
Limited effects of KD on deficits in USV bout structure in juvenile and adult BTBR mice. **(A-H)** Quantification of duration of bouts, total number of calls within bouts, total number of sequences within bouts, and total number of bouts in 5-minute recordings in juvenile **(A, C, E, G)** and adult **(B, D, F, H)** BTBR or B6 mice fed with either SD (circles) or KD (triangles). (**A, B**) Duration of bouts, (**C, D**) total number of calls in a bout, and (**E, F**) total number of sequences within bouts were significantly lower in BTBR than B6 mice. KD treatment showed no effect except for (**C**) an increased number of calls within bouts in juvenile KD-fed BTBR and B6 mice. (**G**) No effect of strain or diet on number of bouts in juveniles. (**H**) Adult BTBR mice displayed a higher number of bouts than B6 mice. There was no effect of KD. Juveniles, B6_SD (*n*=9), B6_KD (*n*=11), BTBR_SD (*n*=9), and BTBR_KD (*n*=11); adults, B6_SD (*n*=10), B6_KD (*n*=10), BTBR_SD (*n*=12), and BTBR_KD (*n*=12). Data expressed as **(E, G)** mean (bars) ± StdDev (error bars) and individual animals (symbols) or **(A-D, F, H)** median (bars) ± IQR (error bars) and individual animals (symbols). *p* values, *** *p* < 0.0001 and * *p* < 0.05.

In summary, these data indicate that – while effects of KD on call bouts were limited – the call bout structure might be slightly more malleable by KD in juvenile than in adult mice, and that adult BTBR mice emit more isolated calls.

### 3.3. KD improved aspects of the composition of call categories

Mouse USVs are remarkably complex and are composed of many different categories distinguishable based on frequency, duration, temporal continuity, and shape of the waveform (Takahashi et al., 2016; Castellucci et al., 2018; Panksepp et al., 2007; Wang et al., 2008; Holy and Guo 2005; Scattoni et al., 2008; Scattoni et al., 2011). These subtypes have been proposed to coordinate context-dependent social interactions (Wright et al., 2010; Burke et al., 2017; Brunelli and Hofer 2007; Hofer et al., 2002). Their utilization has been shown to be altered in a genetic mouse model of autism (Takahashi et al., 2016) and to be malleable including by drugs (Brunelli and Hofer 2007; Simola 2015; Wright et al., 2010). Using VocalMat for MATLAB (Fonseca et al., 2021), we classified the calls we recorded into eleven types (Fig. 5A, B) and further grouped them into three categories (continuous frequency unmodulated, containing frequency jumps, and continuous frequency modulated, Fig. 5C). In either juvenile or adult B6_SD mice, the most common call category contained calls with frequency jumps (∼40% of total calls, Fig. 5C), thereof the step down call type appeared most frequently (19 and 27%, respectively; in BTBR_SD mice: ∼6%, Fig. 5B). In contrast, both juvenile and adult BTBR_SD mice used calls from the unmodulated category most often (64 and 46%, respectively, Fig. 5C), and herein the flat call type (58 and 33%; in B6_SD juvenile and adult mice: 11 and 5%, respectively; Fig. 5B).

**Fig. 5.**
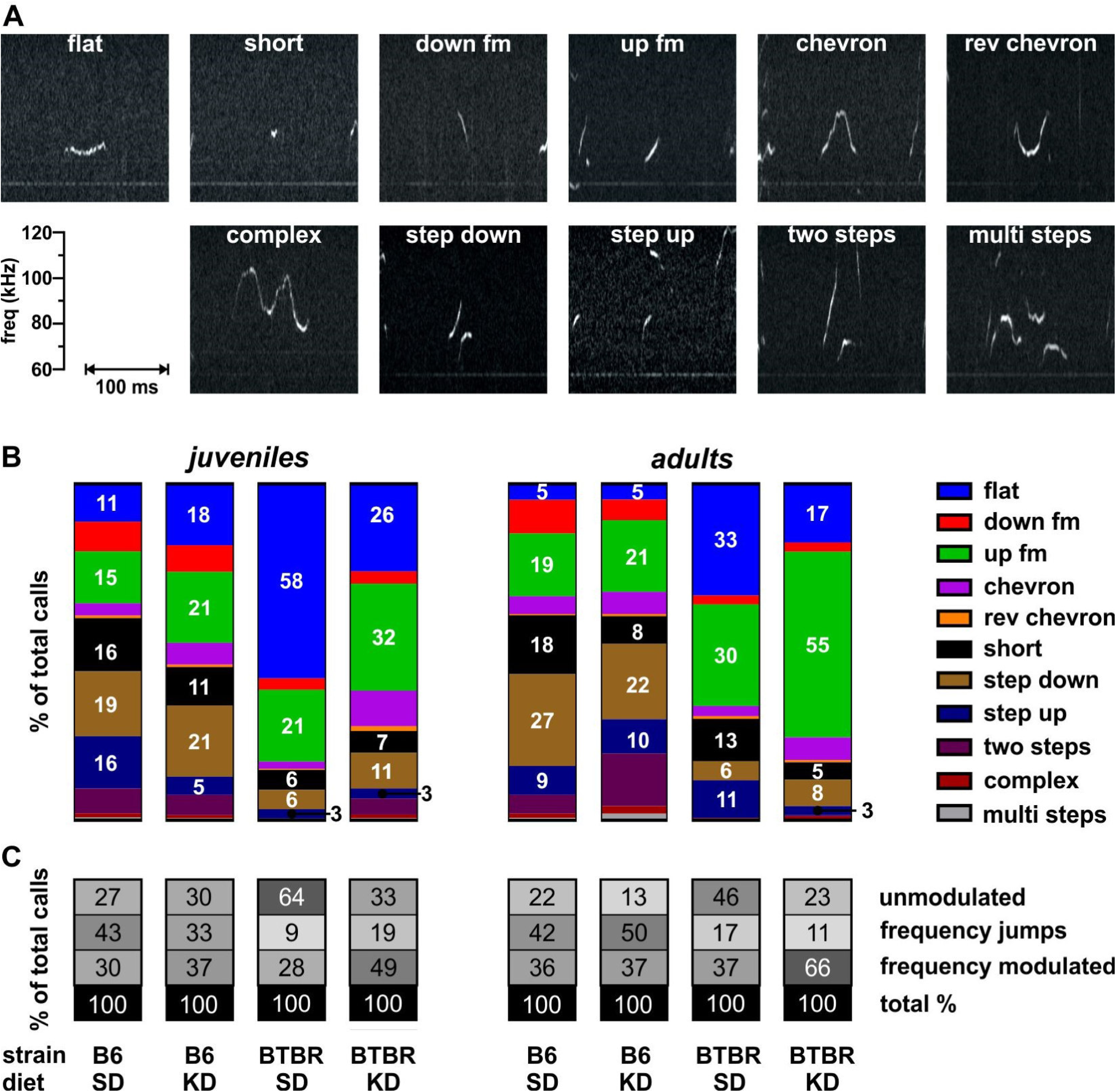
Spectrograms and distributions of the 11 different USV subtypes that the calls were classified into. **(A)** Examples of different call types: flat, short, down frequency modulated (fm), up fm, chevron, reverse (rev) chevron, complex, step down, step up, two steps, and multi steps. **(B)** Call distribution among different call types, shown in different colors, in each experimental group. Number insets denote percent call type. (**C**) Call distribution among different call categories: continuous frequency unmodulated (flat + short), frequency jumps (step down + step up + two steps + multi steps), continuous frequency modulated (down fm + up fm, chevron + rev chevron + complex) in each experimental group. Number insets denote percent call category, darker shades indicate higher percentages. Group information in (**C**) also applies to (**B**). Juveniles, B6_SD (*n*=9), B6_KD (*n*=11), BTBR_SD (*n*=9), and BTBR_KD (*n*=11); adults, B6_SD (*n*=10), B6_KD (*n*=10), BTBR_SD (*n*=12), and BTBR_KD (*n*=12).

Indeed, within the unmodulated call category, the amount of flat call type was significantly higher in both juvenile and adult BTBR_SD mice than in B6_SD or B6_KD mice (Fig. 6A, Ai, Table 3, 4). However, only in juveniles, KD diet significantly decreased flat call usage in BTBR_KD compared with BTBR_SD mice, if not quite to the levels of B6_SD mice (Fig. 6A, Table 3, 4). In adults after KD treatment, BTBR mice still emitted flat calls at a similar level as BTBR_SD mice and significantly more often than B6 mice of either diet (Fig. 6Ai, Table 3, 4). The short call type occurred less often in BTBR mice of both age groups and was not altered through KD treatment (Fig5 F, Fi, Table 3).

**Fig. 6.**
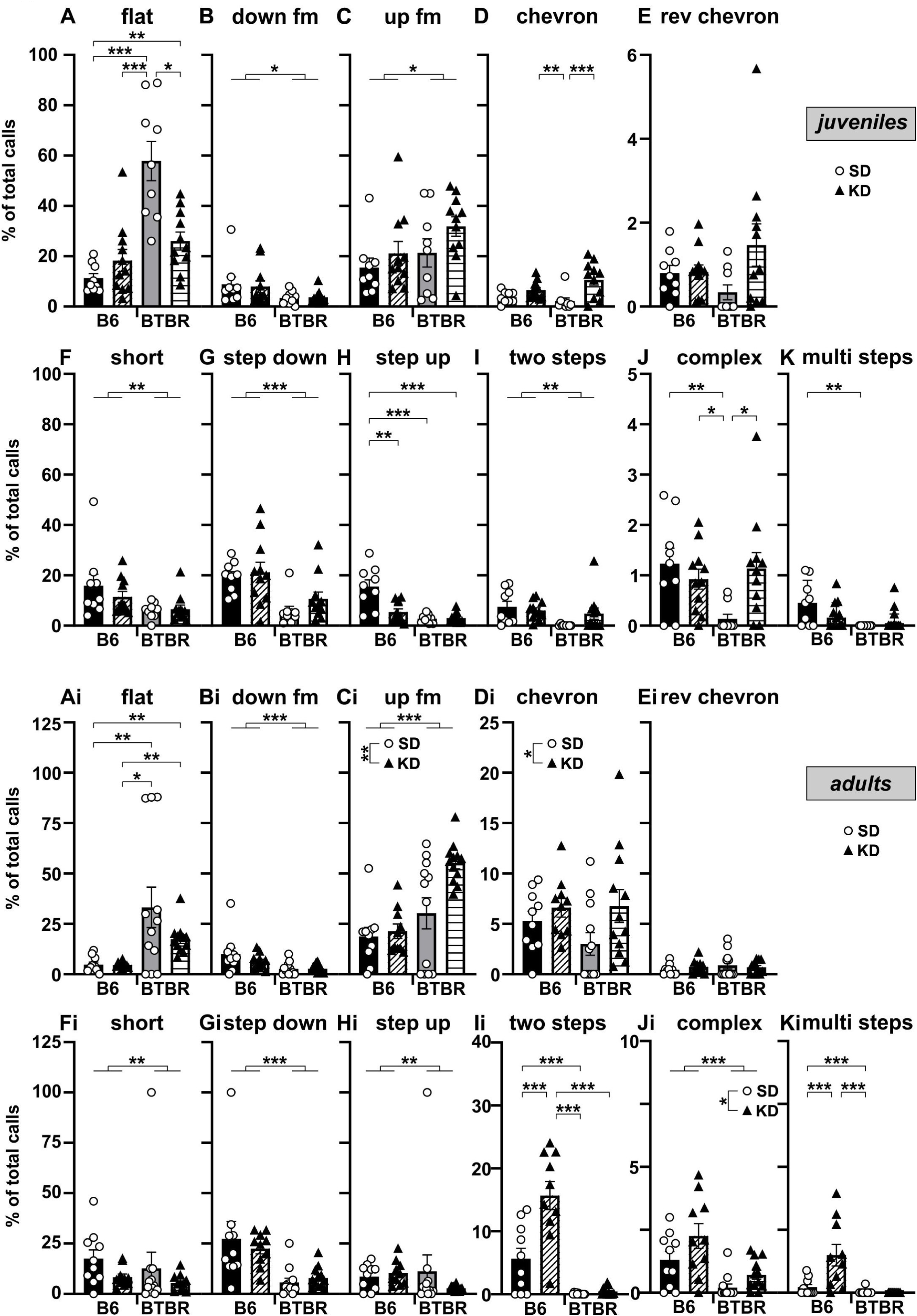
KD improved aspects of the call categories composition more notably in juvenile BTBR mice. Comparison of percentage of total calls by each call type in juvenile **(A-K)** and adult **(Ai-Ki)** BTBR or B6 mice fed with either SD (circles) or KD (triangles). (**A**) Flat: BTBR_KD displayed higher amounts of flat calls compared with B6_SD or B6_KD mice. KD treatment led to a significant decrease in BTBR_KD compared with BTBR_SD mice, however, amounts in BTBR_KD mice were still significantly higher than in B6_SD mice. (**B**) Down fm: Amounts were lower in BTBR mice with no effect of KD diet. (**C**) Up fm: Amounts were higher in BTBR mice with no effect of KD diet. (**D**) Chevron: BTBR_SD were similar to B6_SD mice. KD diet led to a significant increase in BTBR_KD compared with BTBR_SD mice. (**E**) Rev chevron: Amounts were not different between strains or diet. (**F**) Short: Amounts were lower in BTBR mice with no effect of KD diet. (**G**) Step down: Amounts were lower in BTBR mice with no effect of KD diet. (**H**) Step up: Both BTBR_SD or BTBR_KD were significantly lower than B6_SD, with no difference between BTBR mice of either diet. (**I)** Two steps: Amounts were lower in BTBR mice with no effect of KD diet. (**J**) Complex: BTBR-SD displayed fewer complex calls than B6_SD or B6_KD mice. KD diet significantly increased complex calls in BTBR_KD compared with BTBR_SD mice, making them similar to B6_SD or B6_KD levels. (**K**) Multi steps: BTBR_SD mice were significantly lower than B6_SD mice. KD diet did not increase multi step calls in BTBR_KD compared with BTBR_SD mice. (**Ai**) Flat: BTBR_SD mice showed higher amounts than B6_SD and B6_KD mice. Amounts in BTBR_KD were similar to BTBR_SD mice and still higher than in B6_SD and B6_KD mice. (**Bi**) Down fm: Amounts were lower in BTBR mice with no effect of KD diet. (**Ci**) Up fm: BTBR mice used Up fm calls relatively more often than B6 mice. KD diet increased up fm usage in both BTBR and B6 mice. (**Di**) Chevron: BTBR mice were similar to B6 mice. KD diet increased chevron usage in both BTBR and B6 mice. (**Ei**) Rev chevron: Amounts were not different between strains or diet. (**Fi**) Short: Amounts were lower in BTBR mice with no effect of KD diet. (**Gi**) Step down: Amounts were lower in BTBR mice with no effect of KD diet. (**Hi**) Step up: Amounts were lower in BTBR mice with no effect of KD diet. (**Ii)** Two steps: Amounts were lower in BTBR_SD than in B6_SD or B6_KD mice. KD diet significantly increased two step usage in B6_KD compared with B6_SD mice, but not in BTBR_KD compared with BTBR_SD mice. (**Ji**) Complex: BTBR mice showed fewer complex calls than B6 mice. KD diet increased complex call occurrence in both B6 and BTBR mice. (**Ki**) Multi steps: Amounts were lower in BTBR_SD than in B6_SD or B6_KD mice. KD diet significantly increased multi step usage in B6_KD compared with B6_SD mice, but not in BTBR_KD compared with BTBR_SD mice. Juveniles, B6_SD (*n*=9), B6_KD (*n*=11), BTBR_SD (*n*=9), and BTBR_KD (*n*=11); adults, B6_SD (*n*=10), B6_KD (*n*=10), BTBR_SD (*n*=12), and BTBR_KD (*n*=12). Data expressed as mean (bars) ± StdDev (error bars) and individual animals (symbols). Note the differences in Y axes. *p* values, *** *p* < 0.0001, ** *p* < 0.01 and * *p* < 0.05.

**Table 3.**
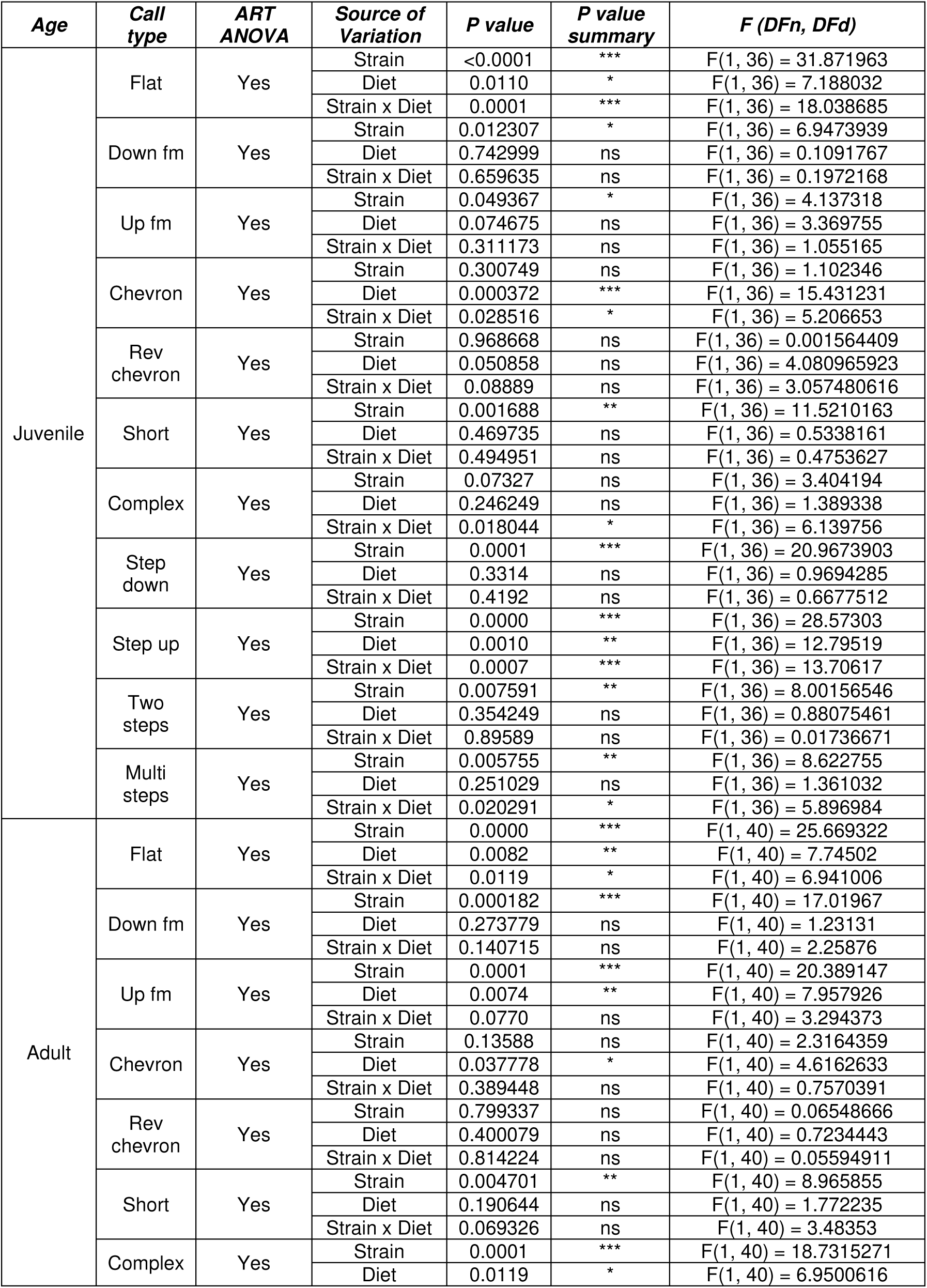
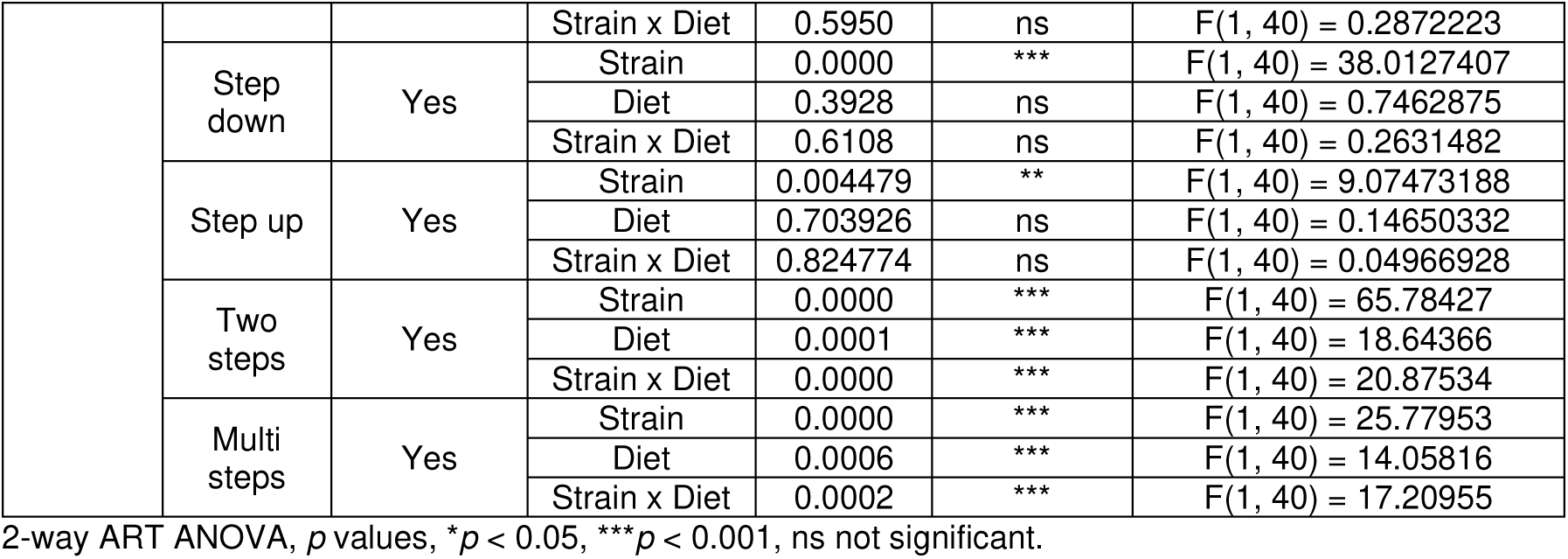
Statistical comparisons of USV call type distribution of calls from mice of the following four groups: juveniles, B6_SD (n=9), B6_KD (n=11), BTBR_SD (n=9), and BTBR_KD (n=11); adults, B6_SD (n=10), B6_KD (n=10), BTBR_SD (n=12), and BTBR_KD (n=12).

**Table 4.** *Post hoc* pairwise comparisons for Strain x Diet interactions of USV call type distribution of calls from mice of the following four groups: juveniles, B6_SD (n=9), B6_KD (n=11), BTBR_SD (n=9), and BTBR_KD (n=11); adults, B6_SD (n=10), B6_KD (n=10), BTBR_SD (n=12), and BTBR_KD (n=12).

| Age | Call type | Contrast | Estimate | SE | DF | t | Adjusted P Value | P value summa |
| --- | --- | --- | --- | --- | --- | --- | --- | --- |
| Juvenile | Flat | BTBR,KD - BTBR,SD | -11.4949 | 3.605713 | 36 | -3.18798 | 0.0150 | * |
|  |  | BTBR,KD - B6,KD | 7 | 3.42068 | 36 | 2.046377 | 0.1904 | ns |
|  |  | BTBR,KD - B6,SD | 12.83838 | 3.605713 | 36 | 3.560567 | 0.0056 | ** |
|  |  | BTBR,SD - B6,KD | 18.49495 | 3.605713 | 36 | 5.129346 | 0.0001 | *** |
|  |  | BTBR,SD - B6,SD | 24.33333 | 3.781704 | 36 | 6.434489 | 0.0000 | *** |
|  |  | B6,KD - B6,SD | 5.838384 | 3.605713 | 36 | 1.619204 | 0.3810 | ns |
|  | Chevron | BTBR,KD - BTBR,SD | 18.37879 | 4.343845 | 36 | 4.230995 | 0.000846 | *** |
|  |  | BTBR,KD - B6,KD | 3.5 | 4.120934 | 36 | 0.849322 | 0.830498 | ns |
|  |  | BTBR,KD - B6,SD | 10.87879 | 4.343845 | 36 | 2.504414 | 0.076199 | ns |
|  |  | BTBR,SD - B6,KD | -14.8788 | 4.343845 | 36 | -3.42526 | 0.008069 | ** |
|  |  | BTBR,SD - B6,SD | -7.5 | 4.555864 | 36 | -1.64623 | 0.366472 | ns |
|  |  | B6,KD - B6,SD | 7.378788 | 4.343845 | 36 | 1.698676 | 0.339218 | ns |
|  | Comple x | BTBR,KD - BTBR,SD | 14.76263 | 4.512589 | 36 | 3.271432 | 0.012097 | * |
|  |  | BTBR,KD - B6,KD | 1.227273 | 4.281017 | 36 | 0.286678 | 0.991645 | ns |
|  |  | BTBR,KD - B6,SD | -1.51515 | 4.512589 | 36 | -0.33576 | 0.986741 | ns |
|  |  | BTBR,SD - B6,KD | -13.5354 | 4.512589 | 36 | -2.99947 | 0.024105 | * |
|  |  | BTBR,SD - B6,SD | -16.2778 | 4.732843 | 36 | -3.43932 | 0.007772 | ** |
|  |  | B6,KD - B6,SD | -2.74242 | 4.512589 | 36 | -0.60773 | 0.928962 | ns |
|  | Step up | BTBR,KD - BTBR,SD | 0.484849 | 4.097077 | 36 | 0.11834 | 0.999396 | ns |
|  |  | BTBR,KD - B6,KD | -5.36364 | 3.886829 | 36 | -1.37995 | 0.519747 | ns |
|  |  | BTBR,KD - B6,SD | -19.1818 | 4.097077 | 36 | -4.68183 | 0.000223 | *** |
|  |  | BTBR,SD - B6,KD | -5.84848 | 4.097077 | 36 | -1.42748 | 0.490967 | ns |
|  |  | BTBR,SD - B6,SD | -19.6667 | 4.297051 | 36 | -4.57678 | 0.000305 | *** |
|  |  | B6,KD - B6,SD | -13.8182 | 4.097077 | 36 | -3.37269 | 0.009277 | ** |
|  | Multi steps | BTBR,KD - BTBR,SD | 5.272727 | 4.072877 | 36 | 1.294595 | 0.572257 | ns |
|  |  | BTBR,KD - B6,KD | -4.90909 | 3.863871 | 36 | -1.27051 | 0.587174 | ns |
|  |  | BTBR,KD - B6,SD | -9.17172 | 4.072877 | 36 | -2.2519 | 0.128781 | ns |
|  |  | BTBR,SD - B6,KD | -10.1818 | 4.072877 | 36 | -2.49991 | 0.076947 | ns |
|  |  | BTBR,SD - B6,SD | -14.4444 | 4.27167 | 36 | -3.38145 | 0.009064 | ** |
|  |  | B6,KD - B6,SD | -4.26263 | 4.072877 | 36 | -1.04659 | 0.723481 | ns |
| adult | Flat | BTBR,KD - BTBR,SD | 2.291667 | 4.1762 | 40 | 0.548745 | 0.94635 | ns |
|  |  | BTBR,KD - B6,KD | 16.4 | 4.380035 | 40 | 3.744262 | 0.003076 | ** |
|  |  | BTBR,KD - B6,SD | 18.25 | 4.380035 | 40 | 4.166633 | 0.000891 | *** |
|  |  | BTBR,SD - B6,KD | 14.10833 | 4.380035 | 40 | 3.221055 | 0.013013 | * |
|  |  | BTBR,SD - B6,SD | 15.95833 | 4.380035 | 40 | 3.643426 | 0.004097 | ** |
|  |  | B6,KD - B6,SD | 1.85 | 4.574797 | 40 | 0.40439 | 0.977336 | ns |
|  | Two steps | BTBR,KD - BTBR,SD | 5.25 | 3.249925 | 40 | 1.615422 | 0.3817 | ns |
|  |  | BTBR,KD - B6,KD | -20.7667 | 3.40855 | 40 | -6.09252 | 0.0000 | *** |
|  |  | BTBR,KD - B6,SD | -8.26667 | 3.40855 | 40 | -2.42527 | 0.0885 | ns |
|  |  | BTBR,SD - B6,KD | -26.0167 | 3.40855 | 40 | -7.63277 | 0.0000 | *** |
|  |  | BTBR,SD - B6,SD | -13.5167 | 3.40855 | 40 | -3.96552 | 0.0016 | ** |
|  |  | B6,KD - B6,SD | 12.5 | 3.560115 | 40 | 3.511123 | 0.0059 | ** |
|  | Multi steps | BTBR,KD - BTBR,SD | -1.54167 | 2.84748 | 40 | -0.54141 | 0.9483 | ns |
|  |  | BTBR,KD - B6,KD | -20.95 | 2.986462 | 40 | -7.01499 | 0 | *** |
|  |  | BTBR,KD - B6,SD | -8 | 2.986462 | 40 | -2.67875 | 0.0502 | ns |
|  |  | BTBR,SD - B6,KD | -19.4083 | 2.986462 | 40 | -6.49877 | 0 | *** |
|  |  | BTBR,SD - B6,SD | -6.45833 | 2.986462 | 40 | -2.16254 | 0.1514 | ns |
|  |  | B6,KD - B6,SD | 12.95 | 3.119258 | 40 | 4.151628 | 0.0009 | *** |

ART-C Tukey's multiple comparisons test, $p$ values, \* $p < 0.05$ , \*\* $p < 0.01$ , \*\*\* $p < 0.001$ , ns not significant.

Within the frequency jump category and in both age groups, KD diet was unable to significantly increase the lower amounts of all call types in BTBR mice, including step down (Fig. 6G, Gi), step up (Fig. 6H, Hi, Table 3, 4), two steps (Fig. 6I, Ii, Table 3, 4), and multi steps call types (Fig. 6K, Ki Table 3, 4).

Furthermore, in the continuous frequency modulated category, KD diet significantly increased the lower usage of complex calls in juvenile BTBR_KD compared with BTBR_SD mice, thereby normalizing the amounts to B6_SD and B6_KD levels (Fig. 6J, Table 3, 4). In adults, the amount of complex call type was also lower in BTBR mice, and increased through KD in both BTBR and B6 mice (Fig. 6Ji, Table 3). In both age groups, KD diet was unable to significantly increase the lower amounts of down fm call type (Fig. 6B, Bi, Table 3). Interestingly, up fm calls were more common in juvenile and adult BTBR compared with B6 mice (Fig. 6C, Ci, Table 3). While KD had no effect on up fm calls in juveniles, it led to a further increase in this discrepancy in adults (Fig. 6Ci, Table 3). The chevron call type was initially not different between the two strains in either age group, but increased through KD diet in juvenile BTBR_KD compared with BTBR_SD (Fig. 6D, Table 3, 4), as well as in adults in both BTBR and B6 mice (Fig. 6Di, Table 3). The rev chevron call type was affected neither by strain nor diet in either age group (Fig. 6E, Ei, Table 3).

In summary, both juvenile and adult BTBR mice showed a different USV call type profile than age-matched B6 mice, most notably less usage of calls containing frequency jumps, and more usage of flat call type. While KD diet reduced the high occurrence of flat call type in juvenile BTBR mice, it failed to do so in adults. In addition, KD diet was unable to improve any call type usage within the frequency jump category in both age groups. Strikingly, within the continuous frequency modulated category, the effect of KD treatment showed a distinct dichotomy between call types and age groups. While KD diet normalized the unusually low amount of the complex call type in both age groups, it exacerbated the excess usage of up fm calls only in adults but not juveniles, and left the rev chevron call type unchanged in both ages. These results indicate that the malleability of call category composition in BTBR mice through KD is dependent on both the age of the animals and the call type. Taken together, in the present study, it appears that KD treatment is more effective to improve call category composition in juvenile BTBR mice. While KD rectified certain call type usage in adults as well, it failed to do so more often than in juveniles or even exacerbated atypically high occurrence of up fm call type.

### 3.4. KD normalized transitions between call types in juvenile, but not adult, BTBR mice

It has been shown that mouse USV call types are not selected in a random order, but rather display a characteristic syllabic structure and are organized into phrases and motifs (Holy and Guo 2005). Using DeepSqueak’s syntax analysis tool (Coffey et al., 2019), we investigated how the syntax flow between call types differed between BTBR and B6 mice with and without KD diet, based on the conditional probabilities of transitions between individual call types within USV bouts.

In juvenile B6 mice with SD diet, the syntax flow path showed transitions between 9 different call types (complex, chevron, up fm, two steps, step up, step down, short, flat, down fm; remaining two call types had frequencies below 0.01 and were therefore excluded from syntax flow path, Fig. 7A). There was a high amount of repetitive use of *e.g*., flat and step down calls, as well as transitions to step down from other call types (such as up fm, two steps, and chevron, Fig. 7A). In BTBR mice with SD diet, the syntax flow path appeared more limited, with only 7 call types (chevron, up fm, step up, step down, short, flat, down fm; remaining call types had frequencies below 0.01 and were therefore excluded from syntax flow path, Fig. 7C). BTBR_SD mice showed prominent repetitive transitions between flat or up fm call types, as well as transitions to flat or up fm from other call types (such as short or chevron, respectively, Fig. 7 C). KD diet in B6 mice did not strongly alter the appearance of the syntax flow path, except for lack of the complex call type (Fig. 7B). In contrast to this, KD diet in BTBR mice greatly changed the flow path composition to 10 call types (complex, chevron, up fm, two steps, step up, step down, short, flat, down fm, rev chevron, Fig. 7D), making the transitions between call types more variable and appear notably more similar to B6_SD mice.

**Fig. 7.**
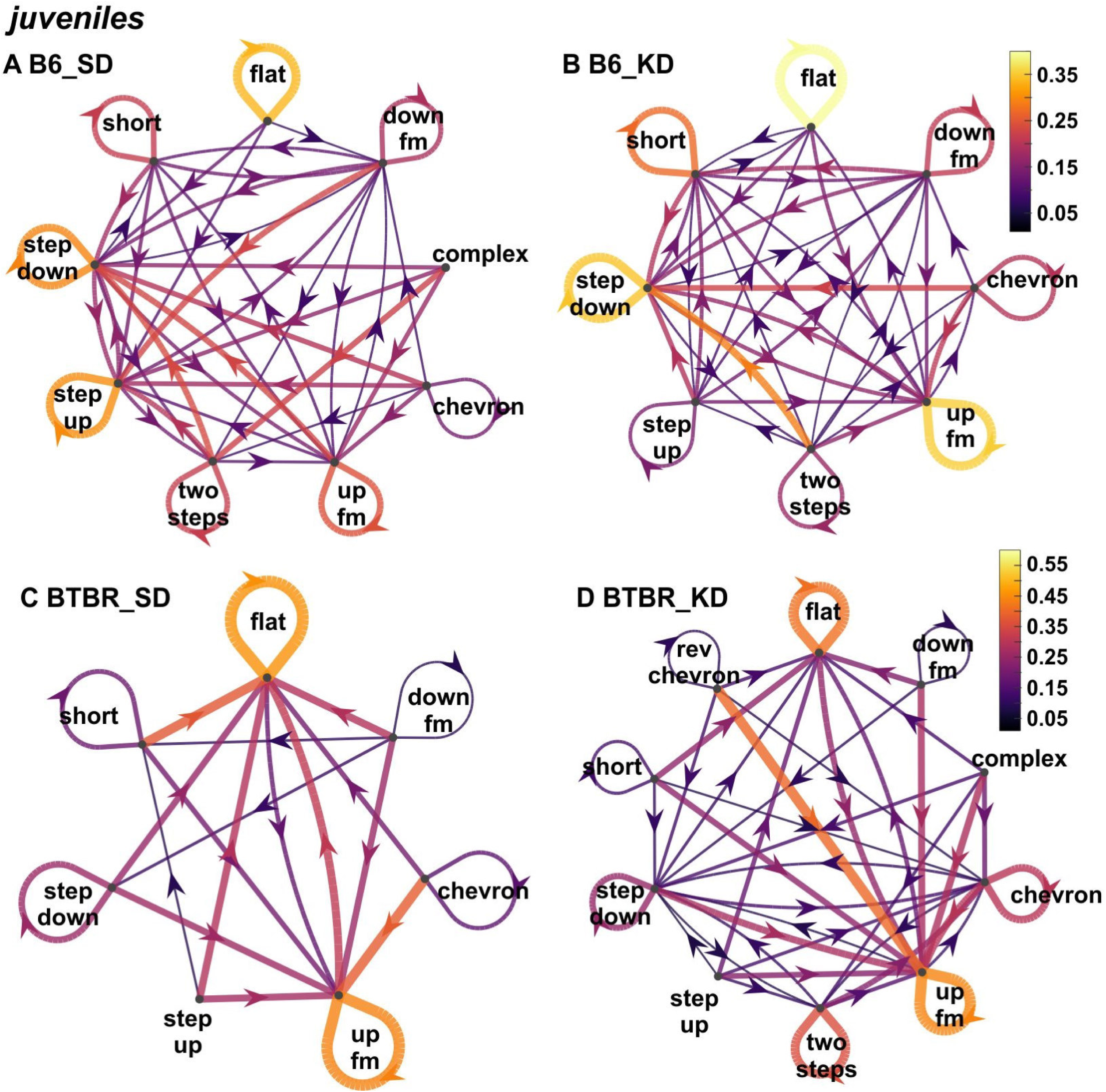
KD improved call type syntax in juvenile BTBR mice. Call type syntax was quantified by means of conditional probabilities for call type transitions within USV bouts in (**A**) B6_SD, (**B**) B6_KD, (**C**) BTBR_SD, and (**D**) BTBR_KD. (**A, B**) B6 mice with either SD or KD diet show diverse call type transitions between 9 or 8 call types, respectively. (**C**) The syntax flow path appeared notably simpler in BTBR_SD mice and included 7 call types. (**D**) KD diet in BTBR mice led to a higher number of call types (*i.e.*, 10) being included in the syntax flow path, which in turn appeared obviously more complex. Arrows represent directions of transitions. Thicker arrows and brighter colours denote higher transition probability. Note the different colour bars for B6 and BTBR mice. B6_SD (*n*=9), B6_KD (*n*=11), BTBR_SD (*n*=9), and BTBR_KD (*n*=11).

Adult B6 mice with SD diet showed transitions between the same 9 call types as their juvenile counterparts (Fig. 8A), with repetitive transitions of step down calls, as well as transitions to step down from other call types (such as two steps and flat, Fig. 8A). The flow path in adult BTBR_SD mice was again more limited, showing transitions between 8 call types (similar to juvenile BTBR_SD plus rev chevron, Fig. 8C) with prominent repetitive transitions between the up fm call type, as well as transitions to up fm from other call types. The syntax flow path in B6 mice after KD treatment appeared quite similar to B6_SD mice, with the exception of additional transitions from multi steps. KD in adult BTBR mice, however, did not broaden the transitioning between call types compared to BTBR_SD mice. Instead, BTBR_KD mice displayed a similarly restricted syntax flow path, with 7 call types (chevron, up fm, step up, step down, short, flat, down fm, Fig. 8D) as well as pronounced transitions between ant to the up fm call type.

**Fig. 8.**
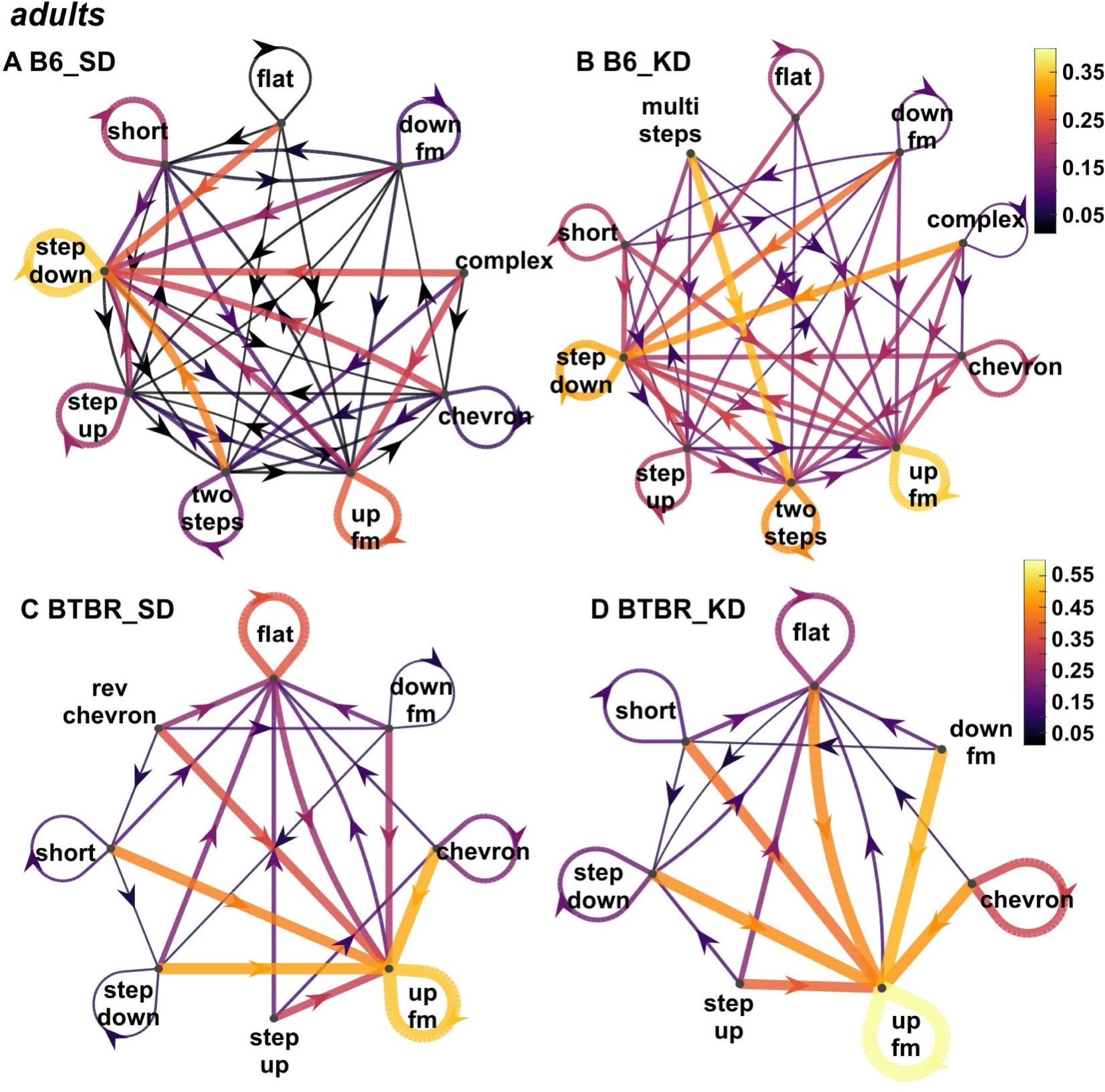
KD did not improve call type syntax in adult BTBR mice. Call type syntax was quantified by means of conditional probabilities for call type transitions within USV bouts in (**A**) B6_SD, (**B**) B6_KD, (**C**) BTBR_SD, and (**D**) BTBR_KD. (**A, B**) B6 mice with either SD or KD diet show diverse call type transitions between 9 or 10 call types, respectively. (**C**) The syntax flow path appeared notably simpler in BTBR_SD mice and included 8 call types. (**D**) The syntax flow path after KD diet contained only 7 call types and restricted transitioning. Arrows represent directions of transitions. Thicker arrows and brighter colours denote higher transition probability. Note the different colour bars for B6 and BTBR mice. B6_SD (*n*=10), B6_KD (*n*=10), BTBR_SD (*n*=12), and BTBR_KD (*n*=12).

In summary, using syntax flow paths based on conditional transition probabilities between individual call types, it seems that both juvenile and adult BTBR mice exert a much simpler syllabic structure than B6 mice. While KD treatment appeared to effectively diversify the syntax flow in juvenile BTBR mice, it was ineffective (or even counterproductive) in adult BTBR mice. This indicates that call type syntax is malleable through KD in juvenile, but not adult, BTBR mice.

## 4. Discussion

### 4.1. Translational approaches for vocal communication in autism

In this study, we report that KD robustly improved ultrasonic vocalization in the BTBR mouse model of autism, not only in the amount and frequency bandwidth, but also in temporal structures of calls as well as call type usage, especially at the juvenile stage. Difficulties in development and usage of language is a hallmark of autism (Stefanatos and Baron 2011; Krishnan et al., 2016; Mayes et al., 2015; Rapin and Dunn 2003), and about thirty percent of children with autism either fail to develop functional language or are minimally verbal (Brignell et al., 2018). Communication is an essential life skill in human society, and studies have found that language deficits are associated with a variety of adverse outcomes, including poorer academic achievement, behavioural difficulties and reduced quality of life. In contrast, functional language usage by school age has been related to better long-term outcomes in autism (Paul 2008). As the underlying mechanisms of these difficulties remain mostly unclear, currently there exist no effective treatments. Behavioral therapies so far constitute the primary intervention, but results have been mixed (Spreckley and Boyd 2009; Paul 2008), especially when minimally verbal children with autism are targeted (Brignell et al., 2018). In addition, these therapies, while highly variable, all require extensive resources from parents and professionals, thus limiting its applicability and reproducibility.

With the huge burden associated with autism (Picardi et al., 2018; Leigh and Du 2015), effective therapies are urgently needed. Studies in both humans and animal models have suggested that KD could be a promising approach for a diverse array of disorders, including neurodevelopmental conditions such as autism, fragile X syndrome, and ADHD (Dai et al., 2017; Boison 2016; Cheng et al., 2017; Gano et al., 2014; Gasior et al., 2006; Lutas and Yellen 2013; Stafstrom and Rho 2012; Yudkoff et al., 2007; Grigolon et al., 2020) as well as in treating refractory epilepsy, especially in children (Neal et al., 2008). This metabolism-based treatment was designed to reproduce the metabolic changes observed during fasting or starvation – adaptations that have been observed historically to render anti-seizure effects. The key metabolic changes include increased blood levels of fatty acids and ketone bodies which are produced by the liver, and decreased glucose. In addition, studies using animal models of diverse diseases have revealed that energy metabolism and mitochondrial function may be relevant targets influenced by KD to induce neuroprotective effects (Gano et al., 2014). Previously, we described that KD diet in juvenile B6 and BTBR mice led to an altered metabolomic profile, including distinct and common pathways in B6 and BTBR mice (Mayengbam et al., 2021). Of the metabolic pathways altered by KD diet (Mayengbam et al., 2021; Heischmann et al., 2018), some are suspected to be involved in atypical social and vocal communication, such as brain region-specific alterations in taurine levels and decreased vocal complexity in a genetic mouse model of autism (Ferreira et al., 2022). However, KD could also act through many other signalling pathways and affect additional molecular targets (Cheng et al., 2017; Stafstrom and Rho 2012; Boison 2016; Gano et al., 2014; Lutas and Yellen 2013; Gasior et al., 2006; Yudkoff et al., 2007; Napoli et al., 2014).

In the present study, our results on juvenile and adult BTBR mice suggest that early treatment could be more effective. Studies in patients of autism also support the notion that early diagnosis and interventions are more likely to have long-term positive effects (Sullivan et al., 2014), although findings in animal models have been mixed. In the case of mouse models of Rett syndrome, restoring expression of the *Mecp2* gene demonstrated that the impairment could be reversed at any developmental stage, from neonatal to adult animals (Guy et al., 2007; Gadalla et al., 2013; Garg et al., 2013). In yet other animal models of autism, early treatment seems to offer better outcomes. For example, in a model of Angelman syndrome, studies have found that introduction of the *Ube3a* allele rescued most syndrome-associated phenotypes early in development, but that the effects were much more restricted later during adolescence (Silva-Santos et al., 2015). In the *Cntnap2* mouse model of autism, chronic treatment with oxytocin during the early postnatal stage led to lasting behavioural recovery (Peñagarikano et al., 2015). In addition, it has been demonstrated that re-expression of *Shank3* gene in adult mice led to improvements in social interaction and repetitive grooming behaviour, while anxiety and motor coordination deficits were not rescued (Mei et al., 2016). Furthermore, a recent study using a rat model of fragile X syndrome showed that pharmacological treatment during adolescent and young adult stage led to sustained correction of deficits in associative learning over several months (Asiminas et al., 2019). Together, these findings suggest that there may be critical developmental windows for optimal treatment in autism.

Other dietary manipulations have also been shown to alter behaviour. For example, the dietary glycemic index, a measure of how carbohydrate content in food affects blood glucose levels, modulates behavioural and biochemical phenotypes in the BTBR mice (Currais et al., 2016). Together, these data support the idea that in the context of genetic predisposition to autism, diet could potentially alter the manifestation of the disease. However, several issues need to be considered. KD is highly restrictive and can be difficult to administer, especially for many people with autism who have meal-related challenges including extremely narrow food selections. In addition, side effects of KD, although most of them are transient and can be managed relatively easily, need to be carefully monitored (Duchowny 2005). As mentioned above, KD was more effective when given to adolescent rather than adult animals. In humans, studies of KD’s effect on development have shown beneficial influences on cognition and behaviour (Pulsifer et al., 2001; Hallbook et al., 2012). However, the impact by KD on development, especially neurodevelopment, needs to be further investigated (Scichilone et al., 2016; Bostock et al., 2017).

### 4.2. Vocalization in mouse models of autism: potentials and limitations

Consistent with previous studies, we found that mouse vocalizations were remarkably complex, in terms of acoustic features of both individual calls and call clusters. Genetically, there are at least several autism risk genes that are involved in human speech and language development, including *Foxp2* and *Cntnap2* (Vargha-Khadem et al., 2005; Haghighatfard et al., 2022; Whitehouse et al., 2011; Rodenas-Cuadrado et al., 2016). Intriguingly, they also play a role in mouse vocalization. In transgenic rodent models that have these genes either deleted or harbouring mutations found in humans with language deficits, altered USV behaviors in either neonatal or adult animals have been observed, including changes in both amount and structure of vocalizations (Peñagarikano et al., 2011; Brunner et al., 2015; Castellucci et al., 2016; Chabout et al., 2016; Schaafsma et al., 2017; Usui et al., 2017; Möhrle et al., 2023). In particular, a reduced vocal repertoire and invariable call sequences with less complicated call types have also been observed in the *Tbx1* and *Cntnap2* mouse model for autism (Takahashi et al., 2016; Burkett et al., 2015) and a higher flat call type usage in poly I:C exposed rat pups (Potasiewicz et al., 2020; Möhrle et al., 2023). These USV alterations are strikingly similar to the findings in BTBR mice in the present study, and might reflect a common feature in mouse models of autism. While it is still a matter of debate into how many functional call categories rodent USVs should be divided, it has been hypothesized that certain call types coordinate moment-by-moment social interactions because of their shared higher relative similarity based on behavioral correlates (Burke et al., 2017). This raises the question if some USV categories might transport specific situational content (Burke et al., 2017; Wright et al., 2010; Panksepp and Burgdorf 2000) and how the call type specific alterations seen in the present study are reflected in on-going behavior influencing a conspecific.

In addition to shared genetic constituents, studies have revealed that anatomical pathways underlying mouse vocalization and human language are also partially conserved. It has been demonstrated that in mice, brain regions involving vocalization include motor cortex and striatal areas, and that the vocal motor cortex sends a direct sparse projection to the brainstem vocal motor neurons, similar to that in humans (Arriaga et al., 2012). Moreover, results from a recent study indicated that the medial preoptic area plays a role in modulating courtship vocalization in mice (Gao et al., 2019). However, a separate study has revealed that mice lacking large parts of the cortex produce mostly normal vocalizations in both pups and adults (Hammerschmidt et al., 2015), suggesting that unlike in humans, cortical structures may not be necessary for the development of mouse vocalizations. In addition, midbrain periaqueductal gray has been recognized as a key region for vocal control in primates and lower mammals, including rodents (Jürgens 2009), and a recent study identified a group of specific neurons in this region that controls social vocalization in mice (Tschida et al., 2019).

Behaviorally, it is generally believed that mouse vocalization is largely innate, thus limiting its application to model development and deficits of human language, which requires the ability to imitate novel sounds (Arriaga and Jarvis 2013). Although unlikely to model all aspects of human speech, mouse vocalization does show plasticity in a context-and experience-dependent manner. Many studies have reported that diverse features of mouse vocalization are modulated by factors such as stage of social (including sexual) interaction, sex and familiarity of the partner during interactions, and the stress level (Yang et al., 2013; Chabout et al., 2015; Grimsley et al., 2016; Hoier et al., 2016; Zala et al., 2017; D’Amato and Moles 2001). These results suggest that although mice have a limited ability to modify vocalization, they could still be used as animal models for understanding certain vocalization features and testing potential interventional approaches in preclinical studies.

### 4.3. Limitations

It is generally believed that mouse vocalization is largely innate, thus limiting its application to model development and deficits of human language, which requires the ability to imitate novel sounds. Therefore, although sharing certain attributes, mouse vocalization is unlikely to model all aspects in the development and deficits of human language.

KD is highly restrictive and can be difficult to administer, especially for many people with autism who have meal-related challenges including narrow food selections. In addition, side effects of KD, although most of them are transient and can be managed relatively easily, need to be carefully monitored. Although some human studies have shown beneficial influences by KD on cognition and behaviour during development, the impact by KD on overall development, especially neurodevelopment, needs to be further investigated. In addition, future studies are required to tease apart the molecular mechanisms of KD’s effects on vocalization.

## 5. Conclusion

This study demonstrated that a brief treatment with KD robustly improved both the amount and bandwidth, as well as temporal and syntax structures of ultrasonic vocalization in the juvenile BTBR mouse model of autism. Together, these results provide further support to the hypothesis that metabolism-based dietary intervention could modify core symptoms in autism. It would be interesting to investigate these pathways in future studies to tease apart the molecular basis of KD, and hopefully provide medicable targets of more specific and potentially personalized approaches.

### Abbreviations

ART: Aligned Rank Transform
B6: C57BL/6J strain
BTBR: Black and Tan Brachyury strain
fm: frequency modulated
ICI: inter-call interval
IQR: interquartile range
KD: ketogenic diet
rev: reverse
SD: standard diet
StdDev: standard deviation
USV: Ultrasonic vocalization

## Acknowledgements

We thank Maryam Khanbabaei for animal husbandry.

## Author contributions

DM participated in MATLAB coding, formal analysis, data interpretation and visualization, and contributed to writing of the original draft. KM participated in data analysis and interpretation, and contributed to drafting the manuscript. JMR provided funding and resources, and helped to design the study. NC designed the study, participated in experiments and data analysis, and contributed to drafting the manuscript. All authors read and approved the final manuscript.

## Data availability

Data will be made available on request.

Appendix A. Supplementary data

