## Supplement Figure 1 and 2 for "Improving vocal communication with a ketogenic diet in a mouse model of autism"

### *Supplementary Material*

#### 1 Supplementary Figures

### Pico-Vac® Mouse Diet 20 5062*

##### DESCRIPTION

Pico-Vac® Mouse Diet 20 is a Constant Nutrition® formulation providing 20% protein for mouse colonies that require extra levels of energy needed for maximum production in post-partum breeding. It is vacuum packed in plastic bags that are irradiated to provide a virtually bacteria-free ration. The vacuum package provides a visual confirmation that the seal on the bag has not been broken.

###### *Features and Benefits*

- Formulated with 20% protein for mouse breeding colonies
- Irradiation gives reliable microbial control and eliminates the need for autoclaving
- Precision processing and selection of highest quality ingredients assures Constant Nutrition® quality
- Designed to meet the energy needs of breeding mouse colonies, transgenic strains, and mice exposed to higher stress levels
- Vacuum packaged in small quantities (2.3 kg/5 lb) for ease of handling and to provide a visual check for package integrity

###### *Product Forms Available*

- Oval pellet, 10 mm x 16 mm x 25 mm length (3/8"x5/8"x1")
- Meal (ground pellets), special order

###### *Other Versions Available*

- 5058 PicoLab® Mouse Diet 20

##### GUARANTEED ANALYSIS

| | |
|---|---|
| Crude protein not less than | 20.0% |
| Crude fat not less than | 9.0% |
| Crude fiber not more than | 4.0% |
| Ash not more than | 6.5% |
| Added minerals not more than | 2.5% |

##### INGREDIENTS

Ground wheat, ground corn, dehulled soybean meal, wheat germ, fish meal, brewers dried yeast, corn gluten meal, porcine animal fat preserved with BHA, soybean oil, calcium carbonate, salt, dicalcium phosphate, monocalcium phosphate, choline chloride, menadione dimethylpyrimidinol bisulfite, DL-methionine, vitamin A acetate, cholecalciferol, pyridoxine hydrochloride, dried whey, folic acid, dl-alpha tocopheryl acetate, biotin, thiamin mononitrate, calcium pantothenate, lecithin, riboflavin, nicotinic acid, casein, vitamin $B_{12}$ supplement, manganous oxide, zinc oxide, ferrous carbonate, copper sulfate, zinc sulfate, calcium iodate, cobalt carbonate, sodium selenite.

##### FEEDING DIRECTIONS

Feed ad libitum to mice. Plenty of fresh, clean water should be available to the animals at all times.

***Mice***-Adult mice will eat up to 5 grams of pelleted ration daily. Some of the larger strains may eat as much as 8 grams per day per animal. Feed should be available on a free choice basis in wire feeders above the floor of the cage.

##### CHEMICAL COMPOSITION[1]

###### *Nutrients*[2]

| | |
|---|---|
| **Protein, %** | **21.8** |
| Arginine, % | 1.15 |
| Cystine, % | 0.31 |
| Glycine, % | 0.93 |
| Histidine, % | 0.50 |
| Isoleucine, % | 1.02 |
| Leucine, % | 1.82 |
| Lysine, % | 1.13 |
| Methionine, % | 0.67 |
| Phenylalanine, % | 0.97 |
| Tyrosine, % | 0.64 |
| Threonine, % | 0.79 |
| Tryptophan, % | 0.25 |
| Valine, % | 1.03 |
| Serine, % | 1.07 |
| Aspartic Acid, % | 2.13 |
| Glutamic Acid, % | 4.47 |
| Alanine, % | 1.34 |
| Proline, % | 1.54 |
| Taurine, % | 0.02 |
| **Fat (ether extract), %** | **9.0** |
| **Fat (acid hydrolysis), %** | **9.1** |
| Cholesterol, ppm | 200 |
| Linoleic Acid, % | 2.32 |
| Linolenic Acid, % | 0.21 |
| Arachidonic Acid, % | 0.02 |
| Omega-3 Fatty Acids, % | 0.32 |
| Total Saturated Fatty Acids, % | 2.72 |
| Total Monounsaturated Fatty Acids, % | 2.88 |
| **Fiber (Crude), %** | **2.2** |
| Neutral Detergent Fiber[3], % | 10.8 |
| Acid Detergent Fiber[4], % | 3.0 |
| **Nitrogen-Free Extract (by difference), %** | **51.8** |
| Starch, % | 39.3 |
| Glucose, % | 0.16 |
| Fructose, % | 0.16 |
| Sucrose, % | 0.71 |
| Lactose, % | 0.78 |
| **Total Digestible Nutrients,%** | **85.3** |
| **Gross Energy, kcal/gm** | **4.60** |
| **Physiological Fuel Value[5], kcal/gm** | **3.75** |
| **Metabolizable Energy, kcal/gm** | **3.56** |

###### *Minerals*

| | |
|---|---|
| **Ash, %** | **5.0** |
| Calcium, % | 0.81 |
| Phosphorus, % | 0.60 |
| Phosphorus (non-phytate), % | 0.33 |
| Potassium, % | 0.70 |
| Magnesium, % | 0.16 |
| Sulfur, % | 0.27 |
| Sodium, % | 0.25 |
| Chlorine, % | 0.42 |
| Fluorine, ppm | 12 |
| Iron, ppm | 200 |
| Zinc, ppm | 120 |
| Manganese, ppm | 120 |
| Copper, ppm | 17 |
| Cobalt, ppm | 0.55 |
| Iodine, ppm | 1.5 |
| Chromium, ppm | 0.56 |
| Selenium, ppm | 0.30 |

###### *Vitamins*

| | |
|---|---|
| Carotene, ppm | Trace |
| Vitamin K (as menadione),ppm | 3.1 |
| Thiamin Hydrochloride, ppm | 15 |
| Riboflavin, ppm | 8.0 |
| Niacin, ppm | 90 |
| Pantothenic Acid, ppm | 21 |
| Choline Chloride, ppm | 2200 |
| Folic Acid, ppm | 2.9 |
| Pyridoxine, ppm | 9.6 |
| Biotin, ppm | 0.30 |
| $B_{12}$, mcg/kg | 51 |
| Vitamin A, IU/gm | 15 |
| Vitamin $D_3$ (added), IU/gm | 3.3 |
| Vitamin E, IU/kg | 57 |
| Ascorbic Acid, mg/gm | — |

###### *Calories provided by:*

| | |
|---|---|
| Protein, % | 23.189 |
| Fat (ether extract), % | 21.635 |
| Carbohydrates, % | 55.176 |

*Product Code

1. Formulation based on calculated values from the latest ingredient analysis information. Since nutrient composition of natural ingredients varies and some nutrient loss will occur due to manufacturing processes, analysis will differ accordingly.
2. Nutrients expressed as percent of ration except where otherwise indicated. Moisture content is assumed to be 10.0% for the purpose of calculations.
3. NDF = approximately cellulose, hemi-cellulose and lignin.
4. ADF = approximately cellulose and lignin.
5. Physiological Fuel Value (kcal/gm) = Sum of decimal fractions of protein, fat and carbohydrate (use Nitrogen Free Extract) x 4,9,4 kcal/gm respectively.

12/11/09

**Supplementary Figure 1. SD nutritional profile.** From: https://www.labsupplytx.com/wp-content/uploads/2012/10/5062.pdf

Bio Serv
Delivering Solutions...
Nutritional • Enrichment • Medicated • Special Needs

### Nutritional Profile

**Product# F3666 -** Ketogenic Diet, AIN-76A Modified, High Fat, Paste, 5 kg/Box

#### Proximate Profile

| | | |
|---|---|---|
| Protein | % | 8.6 |
| Fat | % | 75.1 |
| Fiber | % | 4.8 |
| Ash | % | 3.0 |
| Moisture | % | <10 |
| Carbohydrate | % | 3.2 |

#### Caloric Profile

| | | |
|---|---|---|
| Protein | kcal/gm | 0.34 |
| Fat | kcal/gm | 6.76 |
| Carbohydrate | kcal/gm | 0.13 |
| **Total** | **kcal/gm** | **7.24** |

#### Amino Acids

| | | |
|---|---|---|
| Alanine | gm/kg | 2.3 |
| Arginine | gm/kg | 3.1 |
| Aspartic Acid | gm/kg | 5.5 |
| Cystine | gm/kg | 0.3 |
| Glutamic Acid | gm/kg | 17.3 |
| Glycine | gm/kg | 2.1 |
| Histidine | gm/kg | 2.3 |
| Isoleucine | gm/kg | 4.7 |
| Leucine | gm/kg | 7.1 |
| Lysine | gm/kg | 6.3 |
| Methionine | gm/kg | 2.2 |
| Phenylalanine | gm/kg | 3.8 |
| Proline | gm/kg | 8.7 |
| Serine | gm/kg | 4.8 |
| Threonine | gm/kg | 3.7 |
| Tryptophan | gm/kg | 1.0 |
| Tyrosine | gm/kg | 4.8 |
| Valine | gm/kg | 5.5 |

#### Carbohydrates

| | | |
|---|---|---|
| Monosaccharides | gm/kg | 7.0 |
| Disaccharides | gm/kg | 24.9 |
| Polysaccharides | gm/kg | 0.0 |

#### Fatty Acids

| | | |
|---|---|---|
| C18:2 Linoleic | gm/kg | 115 |
| C18:3 Linolenic | gm/kg | 6.7 |
| Total Saturated | gm/kg | 303 |
| Total Monounsaturated | gm/kg | 288 |
| Total Polyunsaturated | gm/kg | 122 |

#### Minerals

| | | |
|---|---|---|
| Calcium | gm/kg | 5.7 |
| Chloride | gm/kg | 1.7 |
| Copper | mg/kg | 6.6 |
| Chromium | mg/kg | 2.2 |
| Fluoride | mg/kg | 0.0 |
| Iodine | mg/kg | 0.2 |
| Iron | mg/kg | 38.7 |
| Magnesium | gm/kg | 0.56 |
| Manganese | mg/kg | 63.6 |
| Phosphorus | gm/kg | 4.9 |
| Potassium | gm/kg | 3.9 |
| Selenium | mg/kg | 0.19 |
| Sodium | mg/kg | 1128 |
| Sulfur | mg/kg | 366 |
| Zinc | mg/kg | 36.0 |

#### Vitamins

| | | |
|---|---|---|
| Ascorbic Acid | mg/kg | 0.0 |
| Biotin | mg/kg | 0.42 |
| Choline | mg/kg | 274 |
| Folic Acid | mg/kg | 4.2 |
| Niacin | mg/kg | 62.8 |
| Pantothenic Acid | mg/kg | 30.9 |
| Pyridoxine | mg/kg | 12.1 |
| Riboflavin | mg/kg | 12.6 |
| Thiamin | mg/kg | 11.2 |
| Vitamin A | IU/kg | 15500 |
| Vitamin $B_{12}$ | mcg/kg | 21 |
| Vitamin $D_3$ | IU/kg | 2090 |
| Vitamin E | IU/kg | 244 |
| Vitamin $K_3$ (Menadione) | mg/kg | 2.2 |

#### Ingredients

Lard, Butter, Corn Oil, Casein, Cellulose, Mineral Mix, Vitamin Mix, Dextrose

These are typical amounts of nutrients calculated from available information. Actual assay results may vary. For more information contact Jaime Lecker, Ph.D.
908-284-2155 (International)
.

Revised Date: 1/11



Copyright© 2014 Bio-Serv
All Rights Reserved

**Supplementary Figure 2. KD nutritional profile.** From: https://www.bio-serv.com/pdf/F3666.pdf
